# Resolving DJ-1 Glyoxalase Catalysis Using Mix-and-Inject Serial Crystallography at a Synchrotron

**DOI:** 10.1101/2024.07.19.604369

**Authors:** Kara A. Zielinski, Cole Dolamore, Kevin M. Dalton, Nathan Smith, John Termini, Robert Henning, Vukica Srajer, Doeke R. Hekstra, Lois Pollack, Mark A. Wilson

## Abstract

DJ-1 (PARK7) is an intensively studied protein whose cytoprotective activities are dysregulated in multiple diseases. DJ-1 has been reported as having two distinct enzymatic activities in defense against reactive carbonyl species that are difficult to distinguish in conventional biochemical experiments. Here, we establish the mechanism of DJ-1 using a synchrotron-compatible version of mix-and-inject-serial crystallography (MISC), which was previously performed only at XFELs, to directly observe DJ-1 catalysis. We designed and used new diffusive mixers to collect time-resolved Laue diffraction data of DJ-1 catalysis at a pink beam synchrotron beamline. Analysis of structurally similar methylglyoxal-derived intermediates formed through the DJ-1 catalytic cycle shows that the enzyme catalyzes nearly two turnovers in the crystal and defines key aspects of its glyoxalase mechanism. In addition, DJ-1 shows allosteric communication between a distal site at the dimer interface and the active site that changes during catalysis. Our results rule out the widely cited deglycase mechanism for DJ-1 action and provide an explanation for how DJ-1 produces L-lactate with high chiral purity.

## Introduction

Since its discovery in 1997^1^, DJ-1 has been intensively studied owing to its biomedical importance, most prominently its role in certain forms of familial Parkinson’s Disease^2^. Many molecular functions have been attributed to DJ-1, including glyoxalase^3^, deglycase^4^, cyclic 1,3 phosphoglycerate hydrolase^5^, redox sensor^6^, chaperone^7^, RNA binding protein^8^, and others. DJ-1 has a functionally important conserved cysteine residue (Fig. 1a,b) and has long been suspected of being an enzyme, with various activities proposed^3,4,6–9^. One corroborated DJ-1 enzymatic activity is the conversion of methylglyoxal to L-lactate, either directly as a glyoxalase acting on methylglyoxal^3^ or indirectly as a protein/nucleic acid deglycase that repairs early reversible adducts of methylglyoxal and macromolecules^4,10^. Reversibly glycated molecules are in equilibrium with methylglyoxal and thus cannot be easily separated for assay, both activities produce the same products (Fig 1c), and the reaction is difficult to follow spectrophometrically. Therefore, the debate about whether DJ-1 is a glyoxalase or a deglycase has been long-standing and sometimes heated^11–13^, with a growing preponderance of evidence favoring a primary glyoxalase activity^11,14–18^ (Fig. 1d). Disentangling these two activities is complex biochemical problem that requires a direct and conclusive test of these competing hypotheses.

**Figure 1:**
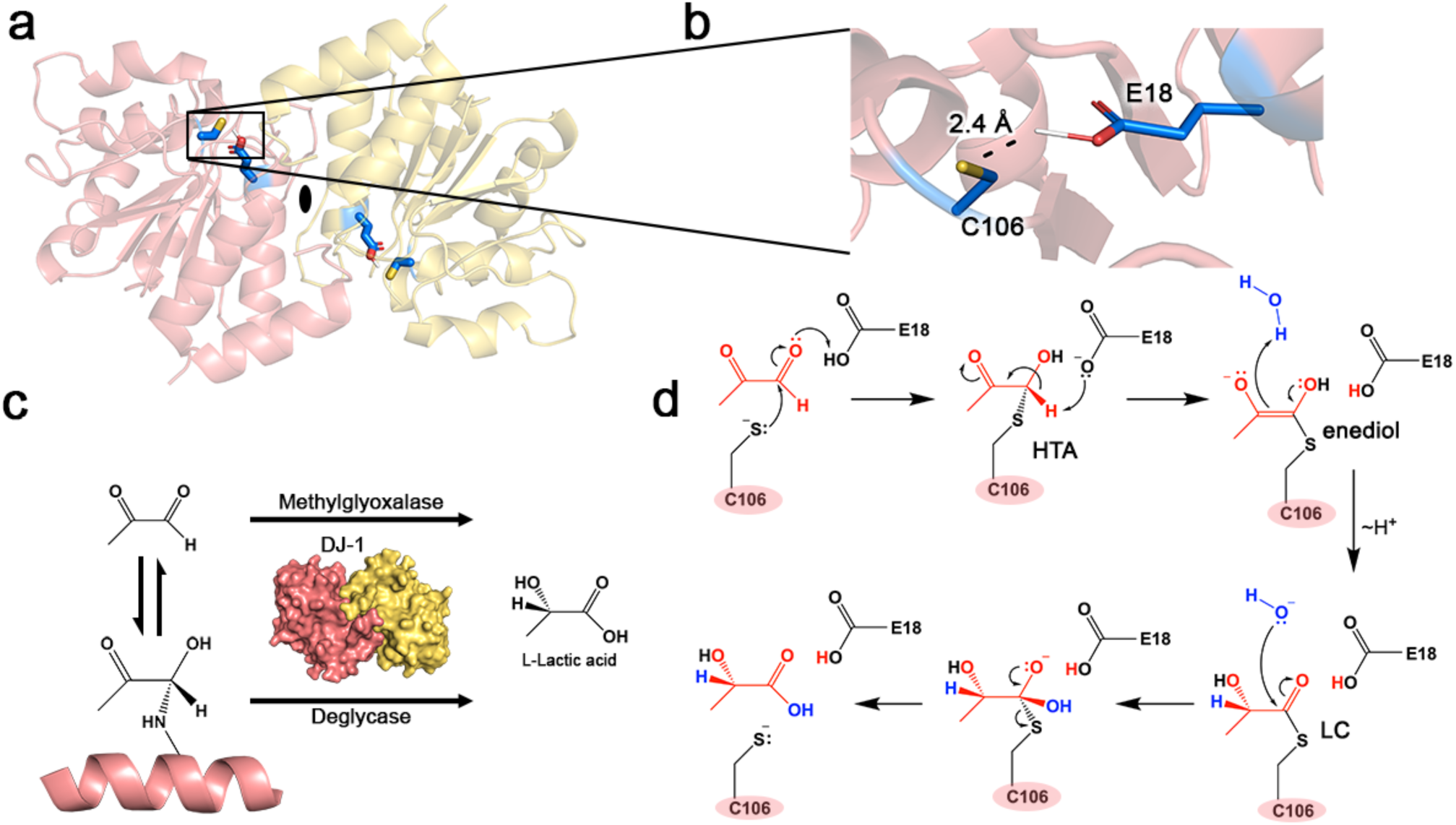
DJ-1 structure, ambiguous substrate, and proposed glyoxalase mechanism. Panel (a) shows the DJ-1 dimer, with protomers colored pink and yellow and the dyad axis of the dimer shown in the black ellipse. The active site centered on C106 and E18 is expanded in panel (b), with the hydrogen bond between the protonated E18 and C106 thiolate shown. Panel (c) illustrates two competing schemes for DJ-1 action, with the top showing direct action on methylglyoxal as substrate and the bottom arrow showing a deglycase activity with modified macromolecules (pink helix) as substrate. Methylglyoxal and reversibly glycated molecules are in equilibrium, shown in the two half-arrows. Both proposed activities produce L-lactic acid. Panel (d) shows a proposed mechanism for DJ-1 glyoxalase activity, with curved arrows showing the direction of electron flow and straight arrows the direction of the reaction. The hemothioacetal (HTA), enediol, and L-lactoylcysteine (LC) intermediates are labeled.

Observing DJ-1 structure during catalysis offers a direct window into its mechanism that would be difficult to obtain using other approaches. Mix-and-inject serial crystallography (MISC) is a powerful technique for answering questions about enzyme mechanisms when the identity of the substrate or catalytic intermediate is uncertain, as direct high resolution structural information can be obtained as the reaction progresses *in crystallo*. MISC relies on the rapid and uniform mixing of microcrystal slurries with a solution of ligand on millisecond timescales followed by delivery of the crystals to the X-ray beam^19,20^, which is technically challenging. Various mixing injectors were initially developed for use at intense X-ray free electron laser (XFEL) sources for time-resolved Serial Femtosecond Crystallography (TR-SFX)^21–24^ and provided detailed views of select enzymatic reactions^25–28^. Despite its exceptional potential to advance the field of structural enzymology, the method has not been widely applied because of its dependence on the availability of limited XFEL beamtime. MISC stands in contrast with other XFEL-pioneered methods, such as serial crystallography, which have been successfully modified for synchrotron sources^29–47^. The growth of synchrotron serial crystallography (SSX) reflects the increased interest in using synchrotrons for these experiments, motivated in part by the more abundant beamtime for structural biology at these facilities. SSX advances have been driven largely by the recent availability of high flux microbeams and fast-framing detectors ^48^, which continue to improve with the advent of fourth-generation synchrotron sources^49^. SSX has been successfully performed on equilibrium, non-evolving species via a variety of sample delivery approaches, including fixed target^31,32,35,37,38,40,43^, tape drive^33,45^, lipidic cubic phase/high-viscosity extrusion^30,34,36,41,47^, and continuous flow^29,39,42^. Although current monochromatic crystallography beamlines at third-generation sources can support SSX using static^31,32,35,37,38,40,43^ or slow moving targets^30,36,41,47,50^, MISC requires a continually flowing sample, which presents challenges related to sample motion during the longer exposures needed at synchrotrons to generate high-quality diffraction data.

Inspired by early time-resolved pump-probe crystallography experiments^51–56^, we turned to Laue crystallography to maximize the X-ray flux in synchrotron MISC experiments. Laue crystallography^57,58^ utilizes a wide X-ray bandwidth (e.g. ∼5% in energy at BioCARS 14-ID beamline at the Advanced Photon Source), which has the advantage of delivering more X-ray flux per exposure. Therefore, a shorter exposure of the microcrystal to the beam is needed to collect usable serial crystallography data, reducing complications arising from crystal motion during X-ray exposure at synchrotron sources. Laue diffraction patterns also capture more information per frame, which reduces sample consumption. The Laue approach has already been applied to SSX experiments^36,44,50,59–62^, and there have even been some TR-SSX studies using light activation^44^. Laue SSX can extend the applicability of time-resolved rapid mixing experiments to the majority of enzymes. However, no such studies have been reported thus far.

Here, we apply this new technology to determine if DJ-1 can catalytically act on methylglyoxal to resolve the long-running dispute about whether it is a glyoxalase or deglycase, and to determine the structure of any potential intermediates at high resolution. The central distinction between these two activities is that a glyoxalase acts directly on methylglyoxal, which is small and easily diffused into a crystal lattice, whereas a deglycase acts on reversibly modified macromolecules that cannot enter the crystal lattice. Therefore, mixing DJ-1 microcrystals with methylglyoxal should result in the formation of key catalytic intermediates in the active site only if DJ-1 is a glyoxalase, while no reaction should occur *in crystallo* if DJ-1 is a macromolecular deglycase (Fig. 1c). Using a Kapton-based flow cell coupled to a microfluidic mixer, we performed pink beam synchrotron MISC at the BioCARS 14-ID beamline (Advanced Photon Source, APS)^63,64^. The use of these new techniques produced time-resolved electron density maps during catalysis, providing direct evidence that DJ-1 is a glyoxalase and allowing it its mechanism to be followed over two cycles of catalysis.

## Results

### DJ-1 cysteine photooxidation and in crystallo modification by methylglyoxal

Past attempts using conventional crystallography with cryotrapping to observe DJ-1 catalytic intermediates failed because introduction of methylglyoxal cracked the 150-300 μm DJ-1 crystals. Microcrystals are often more robust to substrate and ligand introduction than larger crystals^26,65^, and therefore we conducted a serial microcrystallography experiment using ∼25 μm DJ-1 microcrystals (Fig. S1a, Table S1) loaded onto a fixed target sample delivery system^66^ at ID7B2 beamline at the Cornell High Energy Synchrotron Source (CHESS). The microcrystals were illuminated with a monochromatic 12.8 keV (0.6% bandwidth) X-ray beam. Unlike larger crystals, DJ-1 microcrystals survived the introduction of methylglyoxal. Electron density maps for the free enzyme show evidence of oxidation of the active site cysteine (Cys106) to a sulfenic acid (Cys-SOH) (Fig. S1b). This oxidation is observed even when fresh dithiothreitol (DTT) is present, suggesting that it occurs during data collection as a result of X-ray photooxidation^67^. After soaking with methylglyoxal, F_o_(soaked)-F_o_(free) electron density maps show that Cys106 is modified by methylglyoxal, indicating that the substrate enters the crystal and modifies the catalytic nucleophile (Fig S1c). However, it does not establish that DJ-1 can catalyze the reaction (only that it can be modified) and also does not allow information about mechanism be to determined. This result also supports the conclusion that Cys106 oxidation in the free enzyme is due to photooxidation, as an oxidized Cys106 could not react with methylglyoxal and the same batch of crystals were used for both experiments. Therefore, Cys106 oxidation observed in the free enzyme must have occurred subsequent to mixing with substrate and cannot occur when methylglyoxal has already modified Cys106. An important limitation is that, because it takes 13 min to raster through the fixed target chip to collect a full dataset, we could not determine if bona fide catalytic turnover was occurring. We also could not collect structures at timepoints on the seconds timescale that would be needed to probe for distinct intermediates, necessitating a time-resolved experiment.

### Synchrotron mix-and-inject serial crystallography to determine DJ-1 mechanism

To determine crystal structures at defined timepoints after mixing DJ-1 with methylglyoxal, we developed a flow cell that permits MISC to be performed at a synchrotron source. We report here a new implementation of MISC with continuous flow at synchrotrons. To perform these experiments, we coupled our previously developed flow-focused diffusive mixer^21,22^ to a Kapton (Microlumen, Oldsmar, FL) observation region (Fig. S2). Briefly, the central crystal-containing stream is thinned by an external sheath to facilitate rapid diffusion of ligand into protein crystals for reaction initiation (Fig. 2a, Fig. S3). The freshly mixed crystals are then delivered continuously to the X-ray interaction region (Fig. 2a, Fig. S4). The crystal flow speed through the Kapton tube was carefully calibrated to match the residence time of the crystal in the X-ray beam with the exposure time. Kapton has low background scattering and the inner diameter (267 μm) of the tubing was selected to optimize flow speed while reducing background scattering. We utilized this mixer to rapidly combine 50 mM methylglyoxal with DJ-1 crystals to observe the catalytic steps of the reaction. By changing the flowrates and position of the X-ray beam relative to the tip of the mixer, timepoints ranging from 3-30 s could be captured in a single device. To minimize sample consumption, two different mixer designs were used (Fig. S3).

**Figure 2:**
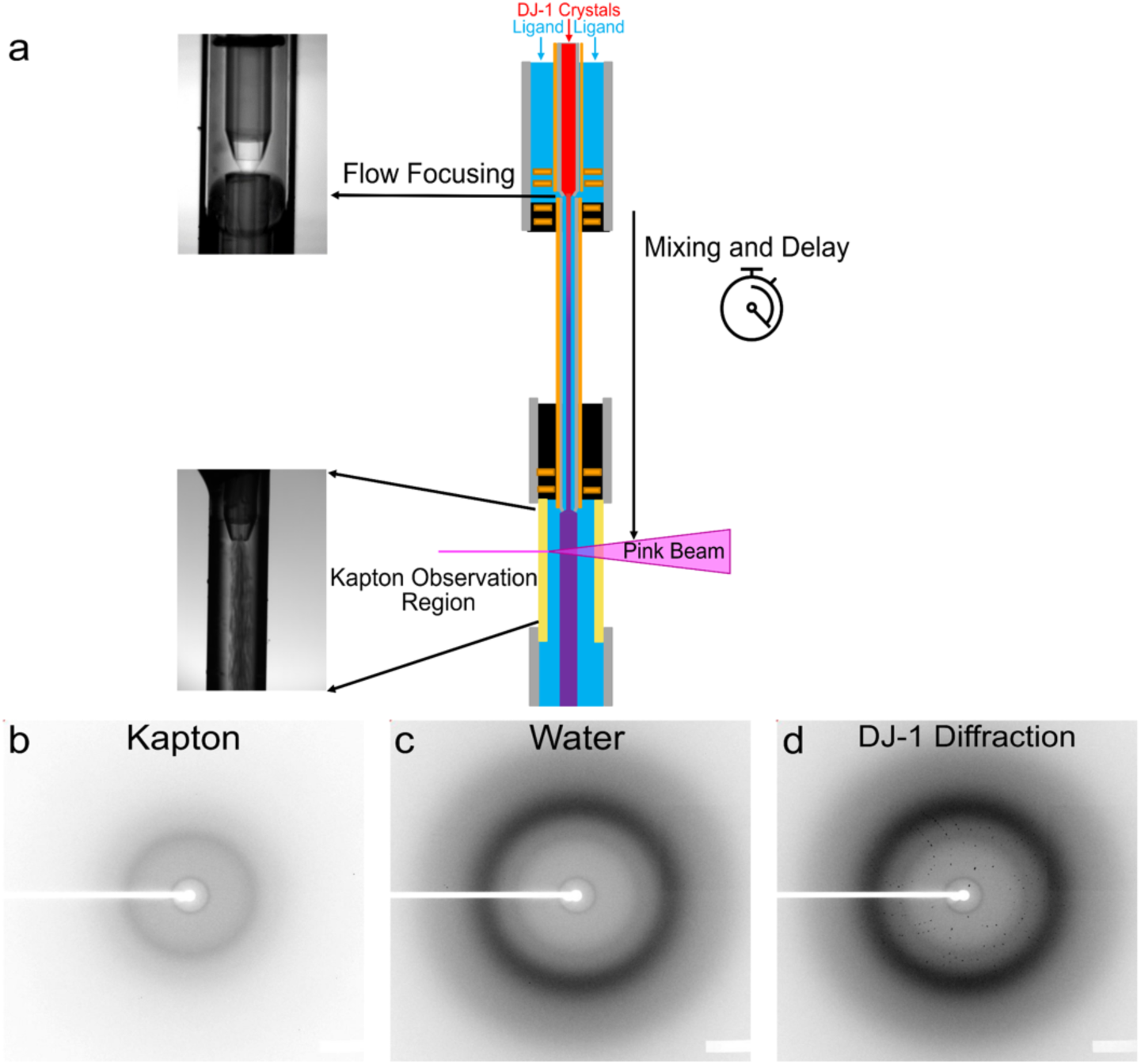
Flow-focused mixing injector for synchrotron MISC applications. Panel (a) shows the flow-focused diffusive mixer is constructed out of concentric capillaries and then coupled directly to a Kapton observation region. The crystal stream (red) is flow-focused by an external ligand stream (blue) to facilitate rapid diffusion of the ligand for reaction initiation. The fully mixed sample (purple) continues to flow and age until a desired timepoint is achieved. Flowrates and the position of the X-ray beam relative to the tip of the mixer can be modified to reach different timepoints. Background scattering from the empty (dry) Kapton tubing (panel (b)), background scattering from water flowing through the Kapton tubing (panel (c)), and representative diffraction pattern from a DJ-1 microcrystal while flowing through the Kapton tubing (panel (d)). Diffraction spots are clearly visible above the background scatter.

Timepoints of 3-30 s were selected to match the slow turnover time of DJ-1, which is approximately 14 sec per reaction (0.07 sec^-1^ turnover number) under these conditions^18^. Timepoint limitations in mixing experiments are primarily dependent on crystal size and the molecular weight and solubility of the ligand. Detailed timepoint calculations are discussed in the Supplemental Information, Table S6. With the high pink-beam flux at the BioCARS 14-ID beamline and the flow cell we used, we found that a crystal size of 25 μm was required for diffraction beyond ∼3 Å when using X-ray exposures no longer than 3.6 µs in order to prevent radiation damage, heating as well as flow cell damage. This size sets the lower limit for the sample stream width for mixing, as the stream width must match the crystal size, influencing the fastest timepoint achievable. With the small methylglyoxal substrate (MW=72.1 Da, D=1.74 x 10^-9^ m^2^/s^39,40^) and a crystal diameter of 25 μm, our theoretical fastest timepoint is 20 ms. At XFELs, our mixers have already been successfully used to capture timepoints on the single ms timescale^27,28^.

All diffraction images were collected using APS 324-bunch mode with an exposure time of 3.6 μs^63^. The background scattering from the Kapton tube used for the observation region with and without water flowing is shown in Fig. 2b-c. A representative diffraction pattern from DJ-1 crystals is shown in Fig. 2d, with diffraction spots clearly visible above the background. All data collection statistics are shown in Table S2. Due to the use of a pink X-ray beam, only 300-1000 Laue diffraction images (hits) were required for a complete dataset for each timepoint. All images were processed using BioCARS’ Python script for hit finding^64^. Then, the hit images were processed by both Precognition (Renz Research, Inc.) and CrystFEL with *pinkIndexer*^60,68,69^ for indexing and integration. The Precognition data were scaled and merged with Careless^70^, while the CrystFEL data continued with CrystFEL pipeline for Laue data processing^60^. We were able to achieve, on average across the timepoints, 0.21 Å better resolution with Careless than with CrystFEL as judged by the canonical CC_1/2_=0.3 resolution cutoff^71^. We attribute this extended resolution primarily to the use of a multivariate Wilson’s prior which models time points as statistically dependent and thereby provides extra constraining information while scaling the reflection observations. This approach has the added advantage of producing higher signal-to-noise, better centered, unweighted time-resolved difference electron density maps. Because of these factors, we present analyses derived from Careless merging results in this manuscript. Nonetheless, we found the dual approach offered complementary advantages, as CrystFEL produces data with resolution-dependent scaling matching the empirical data, while the Bayesian approach used by Careless produces structure factors estimates on a resolution-independent scale^70^. Consequently, the electron density maps calculated from Careless-scaled data have a sharper appearance, similar to what would be expected using B-factor sharpening or the normalized structure factors. By contrast, the results from CrystFEL are more amenable to downstream data analyses which expect observations to match the empirical scale of the data. The multivariate Wilson’s prior implemented in Careless is described in a companion manuscript.

### DJ-1 glyoxalase activity observed using synchrotron Laue MISC

Fixed target serial crystallography of DJ-1 mixed with methylglyoxal provides evidence that the enzyme acts as a glyoxalase and not a deglycase (Fig. S1). To determine the detailed structural mechanism of DJ-1 glyoxalase activity, we collected time-resolved Laue MISC diffraction datasets at 3, 5, 10, 15, 20, and 30 seconds after mixing 25 um DJ-1 microcrystals with 50 mM methylglyoxal. A 0-second timepoint was also collected without mixing. Isomorphous difference (F_o_-F_o_) electron density maps indicate covalent modification of Cys106 at 3 seconds and persisting to the final 30 second timepoint (Fig. 3a-c, S6, S7). Both F_o_-F_o_ and polder omit 2mF_o_-DF_c_ electron density maps show unambiguous evidence of covalent adducts at Cys106 with species derived from methylglyoxal (Fig. 3a-c). The presence of covalent intermediates at Cys106 establishes that methylglyoxal is a direct substrate for DJ-1 and that the protein is not an obligate protein/nucleic acid deglycase.

**Figure 3:**
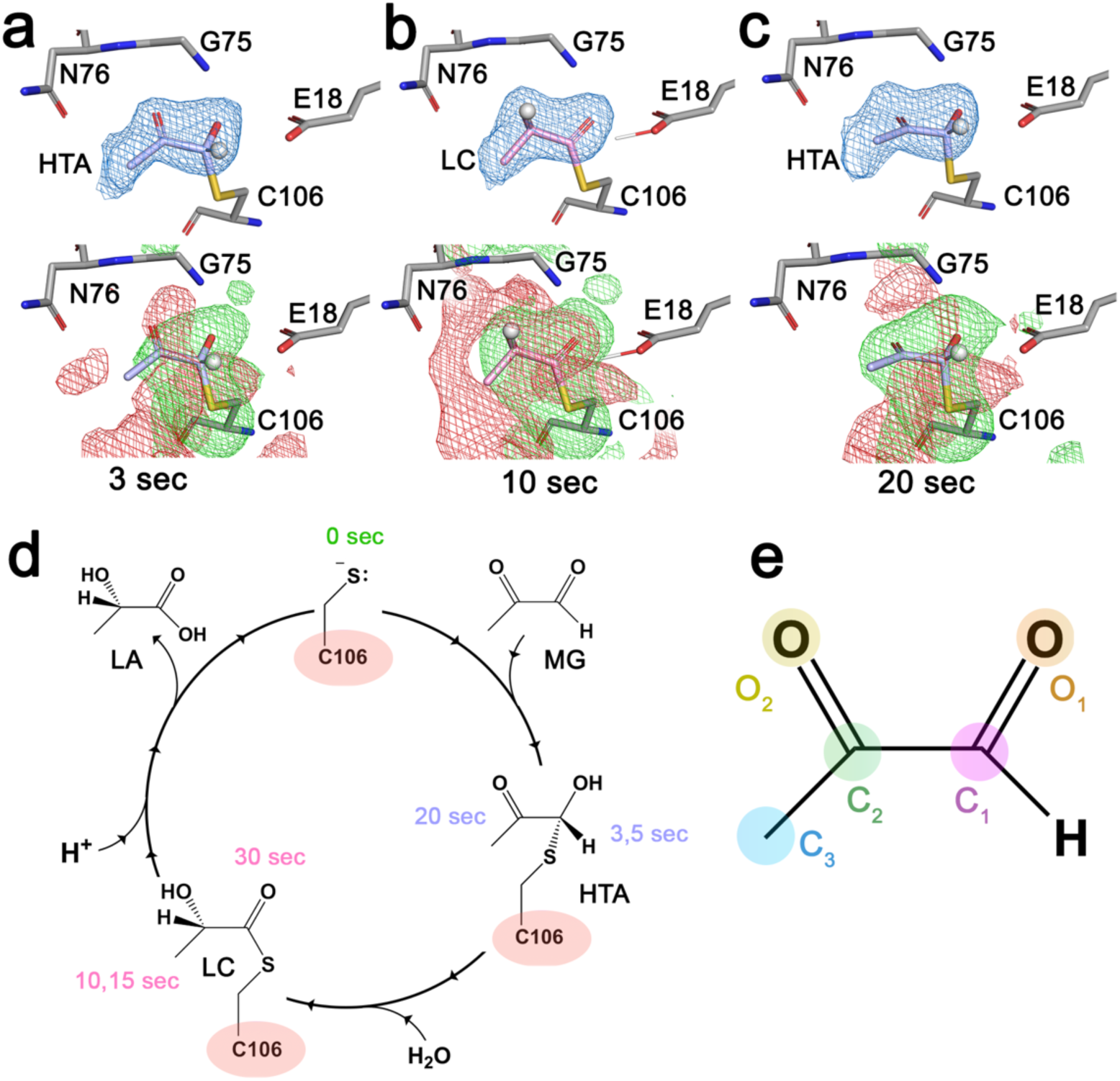
Synchrotron MISC observation of intermediates in DJ-1 glyoxalase catalytic cycle. Panels (a-c) show polder omit electron density at 1 s (blue; top) and F_o_-F_o_ difference electron density at +/- 3σ (green/red, bottom) for the hemithioacetal (HTA, blue bonds)) and L-lactoylcysteine (LC, pink bonds) reaction intermediates. The key hydrogen atom whose location differs in the HTA and LC intermediates is shown as a white sphere. DJ-1 catalyzes two turnovers in the crystal, so the hemithioacetal is observed at 3 and 20 s. Panel (d) shows the catalytic cycle of DJ-1 determined by time-resolved MISC, with the times at which each intermediate is observed shown. Panel (e) shows the non-hydrogen atom number convention for methylglyoxal referenced in the text.

The proposed glyoxalase mechanism of DJ-1 starts with a hemithioacetal adduct, proceeding via an enediol to a lactoylcysteine intermediate which is then hydrolyzed to generate L-lactate (Fig. 1d). The key distinctions between the hemithioacetal, enediol, and lactoylcysteine intermediates are the hybridization of the carbon atoms and the location of protons (Fig. 1d), both of which are difficult to unambiguously determine with these 1.77 Å resolution data. Based on the electron density maps and the mechanism in Fig. 1d, we initially modeled an hemithioacetal adduct for the 3 and 5 second timepoints and an L-lactoylcysteine intermediate for the 10 and 15 second timepoints (Fig. 3a-c). However, the structural similarity of all the proposed intermediates makes determining which species are present difficult based solely on F_o_(ι1t)-F_o_(0s) and polder omit electron density maps. We addressed this by considering a matrix of F_o_-F_o_ electron density maps calculated across all paired timepoints (Fig. S5, S6). F_o_-F_o_ maps are sensitive to small differences between otherwise similar species and thus a full inter-timepoint matrix can disambiguate complex time-series X-ray diffraction data. Based on this F_o_-F_o_ matrix, we conclude that the 3, 5, and 20 second timepoints are the same species (proposed hemithioacetal) as are the 10, 15 and 30 second species (proposed L-lactoylcysteine) (Fig. 3d, Fig. S7). Identical species at different timepoints will not generate strong F_o_-F_o_ electron density, as shown by the F_o_-F_o_ matrix (Fig. S5, S6). This analysis indicates that DJ-1 catalyzes nearly two full turnovers in the crystal in 30 seconds, closely matching expectations based on its solution turnover number of 0.07 sec^-1^ ^14,18^. Inspection of the F_o_-F_o_ maps between the L-lactoylcysteine and hemithioacetal intermediates (e.g. F_o_(10s)-F_o_(3s)) shows that the C_2_, C_3_, and O_2_ atoms (Fig. 3e) become more ordered upon formation of the L-lactoylcysteine (positive F_o_-F_o_ density peaks in Fig. S6, S7), potentially stabilizing this intermediate and helping to drive the proposed 1,2 proton shift required to progress from hemithioacetal to L-lactoylcysteine (Figs. 1d, 3d). The enediol was not modeled, consistent with it being a reactive species with low expected occupancy in the active site, although it is possible that there is a mixture of intermediates at these timepoints and that we have modeled only the predominant one for each timepoint.

### DJ-1 catalytic mechanism and allostery

The time-resolved structures of DJ-1 acting on methylglyoxal allow us to clarify several aspects of its mechanism. The catalytically essential Glu18 residue ^3,18^ is positioned to act as a general acid that protonates the O_1_ atom of the hemithioacetal upon attack of the Cys106 thiolate and is also a candidate for the base that subsequently abstracts the C_1_ proton to form the enediol (Fig. 3a-c). Contrary to some prior proposals ^16,72^, His126 does not appear to play any direct role in DJ-1 catalysis, as F_o_-F_o_ electron density shows that it is pushed away by formation of covalent intermediates (Fig. 4a). These data indicate that the significance of His126 is as a dimer-spanning hydrogen bonding residue ^73,74^ and not as a general acid/base. The position of the hemithioacetal in the DJ-1 active site also provides an explanation for the high enantiopurity of the L-lactate (S-lactate in CIP notation) produced by DJ-1, as the *re*-face of the C_2_ atom is exposed to a nearby ring of ordered solvent molecules, while the *si*-face is occluded by protein contacts (Fig. 4b,c). Therefore, we propose that solvent preferentially protonates the C_2_ atom at the *re*-face of the enediol intermediate, selectively producing L-lactate (Fig. 4d).

**Figure 4:**
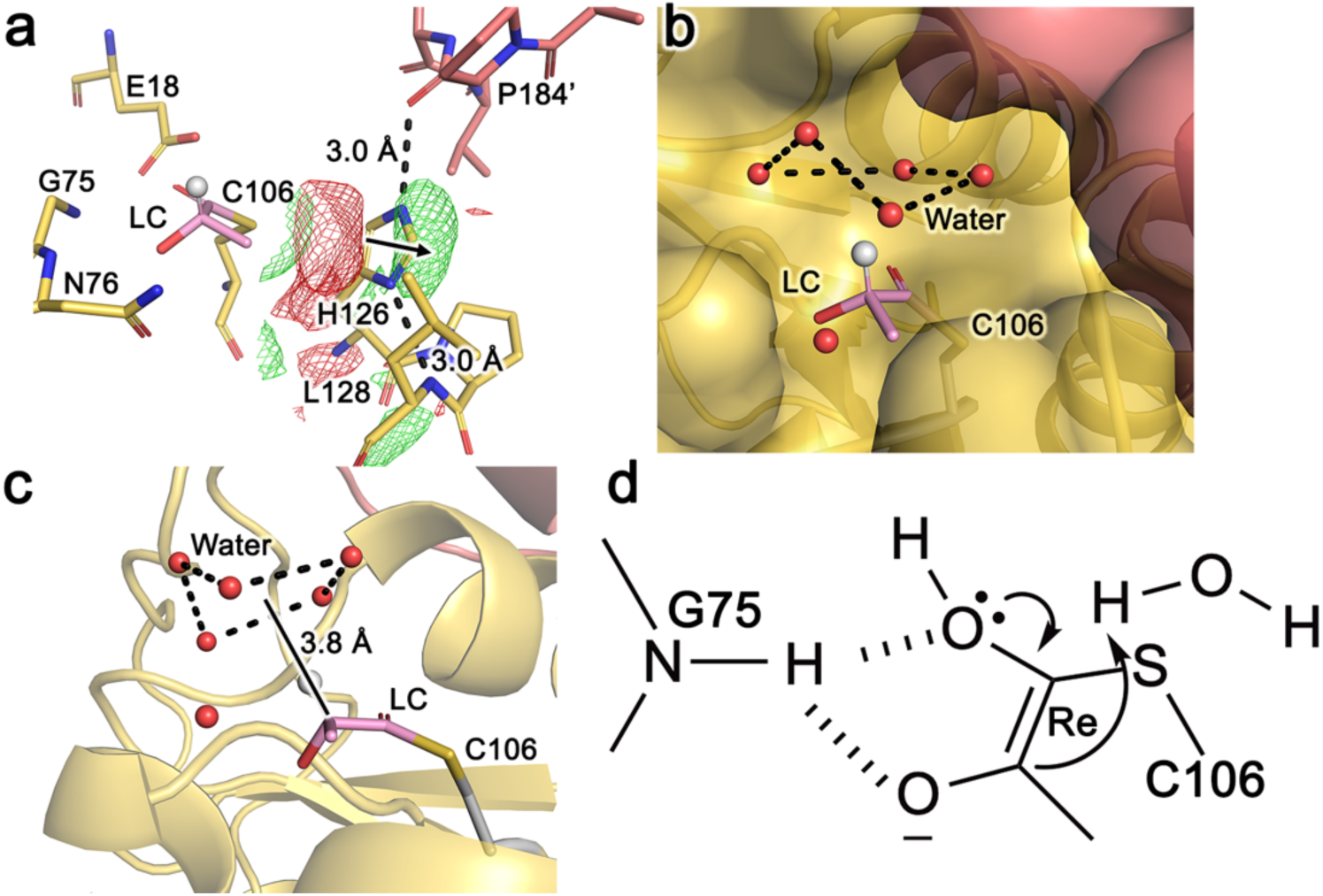
DJ-1 produces L-lactate owing to preferential facial solvent exposure of a reactive intermediate. Panel (a) shows F_o_(10s) -F_o_(0s) difference electron density at +/- 3σ (green/red) for the L-lactoylcysteine intermediate (pink bonds), showing that H126 moves away from the intermediate (black arrow) and is not directly involved in catalysis. Panels (b) and (c) show two views of the active site pocket and a nearby ring of ordered solvent that is the likely source of the proton that sets stereochemistry at the C_2_ atom in L-lactoylcysteine (LC, pink bonds). Panel (d) shows that preferential re-face solvation of the proposed enediol intermediate results in DJ-1 enantiopure L-lactoylcysteine and the final L-lactate product.

DJ-1 does not display features in the F_o_-F_o_ map that indicate widespread correlated motions during catalysis, in contrast to the related enzyme isocyanide hydratase ^26,75^ or the unrelated fluoroacetate dehalogenase^38^. However, F_o_-F_o_ map features near Cys53 change in synchrony with changing intermediates in the active site (Fig. 5, Fig. S8, S9, S10), demonstrating that Cys53 and Cys106 are in allosteric communication. The refined models show a minor wiggle of the Cys53 thiols as DJ-1 proceeds through its reaction cycle, which generates an 8-12σ (0.18-0.31 e^-^/Å^3^) F_o_-F_o_ map signal owing to the high electron density of sulfur (Fig. S10). This demonstrates that sulfur-containing residues have value as sensitive reporters of dynamics in time-resolved crystallography. There are few prominent F_o_-F_o_ density features at 3.0 σ or greater between these two sites, and therefore we cannot identify a candidate pathway for allosteric communication spanning the 24 Å distance between Cys106 and Cys53. Despite this, prior work has indicated that Cys53 can influence Cys106 oxidation^76,77^, providing circumstantial evidence for a physiological connection between these two sites that is consistent with the time-resolved crystallography result.

**Figure 5:**
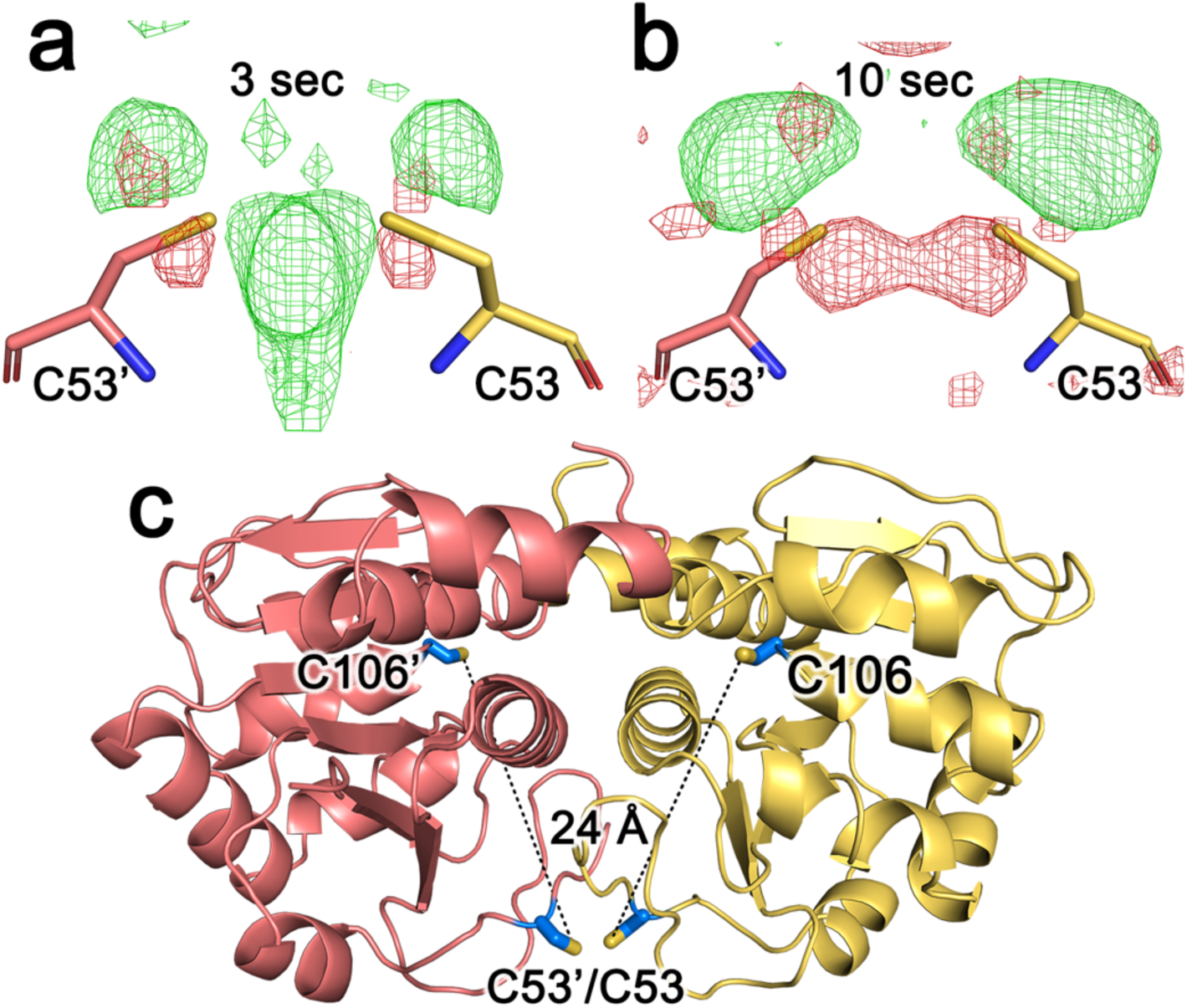
Allosteric communication between active site and distal cysteine residues at the dimer interface during DJ-1 catalysis. Panel (a) shows F_o_ -F_o_ difference electron density maps calculated at the indicated time points at +/- 3σ (green/red) near C53 at the dimer interface. This residue moves during catalysis because it is in allosteric communication with the active site that is 24 Å away (panel (c)).

## Discussion

Our time-resolved synchrotron MISC experiment shows that DJ-1 is a glyoxalase and identifies two primary intermediates as DJ-1 converts methylglyoxal to L-lactate. Both fixed-target and time-resolved serial crystallography establish that DJ-1 does not require a glycated substrate and is not an obligate deglycase^10,12^. These results provide direct structural evidence supporting a growing number of enzymology studies also indicating that DJ-1 is not a deglycase, and thus we join calls^11,14–16,18^ to correct the erroneous attribution of DJ-1 as a “protein/nucleic acid deglycase” that has pervaded the literature and databases.

Time-resolved structures of DJ-1 in the presence of methylglyoxal support the mechanism shown in Fig. 1d. Our data show that the Cys106 thiolate attacks the electrophilic aldehyde carbon atom of methylglyoxal to form a species that we assigned as a tetrahedral hemithioacetal intermediate. This is followed by hydrogen atom rearrangement, either through a 1,2 proton shift (Fig. 1d) or a concerted hydride shift (Fig. S11), two mechanisms between which our data cannot discriminate. We favor the 1,2 proton shift because it is the accepted mechanism of the glutathione-dependent glyoxalases, requiring formation of an enediol intermediate with sp^2^-hybridized C_1_ and C_2_ atoms. Although these two mechanisms have the hemithioacetal and L-lactoylcysteine intermediates in common, they could be tested using isotope retention analysis of methylglyoxal deuterated at the aldehyde hydrogen. Our synchrotron MISC results shows that hemithioacetal intermediate proceeds to L-lactoylcysteine by a highly selective protonation of the solvent exposed *re*-face of the C_2_ atom, while the *si*-face is occluded by protein-intermediate contacts. The final step of catalysis is an attack by water at the C-S bond to release L-lactate. We do not observe an ordered water that is properly positioned for this attack in the electron density, although it is probable that it would have to enter the active site near Glu18 (Fig. 1d).

The physiological relevance of DJ-1’s glyoxalase activity is unresolved. Some prior work, including our own, indicates that DJ-1 can modestly decrease certain methylglyoxal adducts even in the presence of the more active glyoxalase I/II system, although cellular viability is not affected ^18,78^. Some studies have reported that DJ-1 has a larger role in reducing cellular glycation burden and can improve cellular viability during methylglyoxalase challenge ^3,4^, while others indicate no effect^11,14^. An intriguing new activity has been proposed in two recent studies showing that DJ-1 can protect macromolecules and nucleophilic metabolites against modification by a cyclic 1,3 phosphoglycerate metabolite with a k_cat_ that is four orders of magnitude higher than the methyglyoxalase activity^5,79^. This cyclic 1,3-phosphoglycerate hydrolase activity also requires Cys106 and likely involves Glu18 as a general acid/base, suggesting mechanistic similarities with the glyoxalase activity of DJ-1 that could be explored using time-resolved X-ray crystallography in future work.

Our new flowcell demonstrates that MISC can be performed at synchrotrons, which will expand the reach of the technique. The fastest timepoint achievable at synchrotrons will depend on the minimum crystal size needed to obtain high-quality diffraction at a particular beamline, as the crystal size limits the fastest accessible mixing time for a given ligand owing to diffusion. Timepoints of tens of milliseconds and beyond are easily accessible using ∼10 micron crystals, which is a relevant timescale for many biological processes. Synchrotron MISC can also be used to improve success of XFEL beamtimes, because the mixer employed here can, without modification, be coupled to a gas dynamic virtual nozzle and used at an XFEL^4^. Longer MISC timepoints could be collected at synchrotrons in advance of XFEL beamtimes to help determine relevant timescales *in crystallo,* which remains a challenge for many systems that lack spectroscopically active species, such as DJ-1. Faster timepoints could then be captured at XFELs with smaller crystals. Our results demonstrate that synchrotron MISC using pink X-ray beams can resolve difficult questions about enzyme mechanisms and is accessible to a large user community interested in time-resolved structural enzymology.

## Methods

### Expression and purification of DJ-1

Recombinant DJ-1 was expressed similarly to previously described methods^80,81^ with minor changes. DJ-1 cloned into pET-15b (Novagen) was expressed with a thrombin-cleavable N-terminal hexahistidine tag in BL21(DE3) *E. coli* cells grown in Miller LB broth (Fisher Bioreagents BP1426-500) supplemented with 100 μg/ml ampicillin and shaken at 250 rpm at 37°C. Protein expression was induced when the culture reached an OD_600_ = 2 by addition of isopropyl β-D-1-thiogalactopyranoside (IPTG) (GoldBio I2481) at a final concentration of 400 μM. Cells were grown for an additional 6 hours and then pelleted by centrifugation and stored at -80°C.

The cell pellet was resuspended in 10 ml of lysis buffer (50 mM HEPES, 300 mM NaCl, degassed) per gram of pellet. Lysozyme (GoldBio L-040-10) was added to a final concentration of 4 mg/ml and incubated at 4°C for 30 – 60 minutes with gentle stirring. The cell suspension was then sonicated at 4°C using a Misonix Sonicator 3000 and then clarified by centrifugation at 30,000 x g for 30 minutes. The clarified lysate was passed through Ni^2+^-NTA resin (Millipore Sigma cat. no. P6611) and washed with lysis buffer until no protein could be detected in the wash using Bradford’s reagent. DJ-1 was eluted from the column with vacuum-degassed elution buffer (50 mM HEPES pH=7.5, 300 mM NaCl, 250 mM imidazole) and then mixed with thrombin (MP Biomedicals cat. no. 154163) at a ratio of 1.5 U thrombin per 1 mg DJ-1. The protein/thrombin mixture was dialyzed against 25 mM HEPES pH=7.5, 150 mM KCl at 4°C overnight. DJ-1 was then passed through Ni^2+^-NTA (Millipore Sigma cat. no. P6611) to remove any uncleaved protein, followed by benzamidine sepharose 4 resin (Cytiva cat. no. 17512310) to remove thrombin. DJ-1 purity was verified using overloaded Coomassie-stained SDS-PAGE. Purified DJ-1 was then concentrated to 45 mg/mL determined using an extinction coefficient at 280 nm (χ_280_) of 4000 M^-1^cm^-1^, flash frozen in liquid nitrogen in 50-500 μl aliquots, and stored at -80°C.

### DJ-1 microcrystal growth

DJ-1 was crystallized as previously described^80^ using hanging drop vapor equilibration against a reservoir of 100 mM HEPES, 200 mM NaCl, 15% PEG 3350. DJ-1 at 45 mg/mL was mixed in a 1:1 ratio with the reservoir for a total drop volume of 4 μL and incubated at room temperature (∼20°C) until bipyramidal 200-300 μm crystals formed in 2-3 days.

Microcrystals were grown by seeding using pulverized macroscopic crystals as described previously^26^. Approximately 50-100 macrocrystals measuring 200-300 μm in each dimension were harvested into 500 μL of reservoir solution, mixed with approximately 50-100 0.5 mm diameter stainless steel beads (Next Advance cat. no. SSB05) and vortexed for 10 minutes or until most crystals were no longer visible in the microscope. The seed stock solution was centrifuged at 100 x g for one minute to remove any unwanted larger crystal fragments. Aliquots of the seed stock were diluted into 2x stabilizing solution (200 mM HEPES pH=7.5, 400 mM NaCl, and 30% PEG 3350), rapidly mixed with an equal volume of DJ-1 at 45 mg/ml in storage solution, incubated at room temperature for approximately 30 minutes, and then diluted two-fold with 1.15x reservoir solution (115 mM HEPES pH=7.5, 230 mM NaCl, and 17.25% PEG 3350) to stop crystal growth. A 1:150 v:v dilution of seed stock produced uniform 25 μm x 25 μm x 25 μm DJ-1 microcrystals. Before use, DJ-1 crystals were further diluted two-fold and passed through a steel mesh filter with 25 μm pores (Pure Pressure, Denver, CO).

#### Data collection and analysis at ID7B2, CHESS

DJ-1 microcrystals were loaded into Kapton fixed target chips as previously described.^66^ Briefly, ∼5-10 μL of DJ-1 microcrystals were pipetted and added directly to the chip inside a controlled humidity box. Suction from the backside of the chip was used to remove excess supernatant. The chip was sealed with pieces of Mylar film. Then, the chip was mounted directly onto the goniometer at ID7B2, CHESS. The beam size was 20 μm x 20 μm and the X-ray energy was 12.8 keV with a 0.6% bandwidth. The chip was raster scanned to collect data at different positions throughout the chip. At each position, 5 images were taken, with an exposure time of 10 ms per frame and an oscillation of 0.25°.

For the soaking experiments, DJ-1 crystal growth was quenched as described above. Then, the crystals were gently spun down, and the crystals were washed five times to remove DTT. Next, a stock concentration of DJ-1 was added to the crystals to reach a final concentration of 40 mM methylglyoxal and 2 mM DJ-1. The crystals were immediately loaded onto the chip for data collection. All CHESS data was analyzed using DIALS^82^ and custom CHESS scripts as described on Github (https://github.com/FlexXBeamline/ssx-dials).

### Mixing device fabrication

A flow-focused diffusive mixer was constructed out of concentric capillaries and coupled directly to a Kapton observation region, similar to our previous work^21,22,83^. Briefly, a fused silica capillary (75-100 μm ID, 200 μm OD, Polymicro Technologies, Phoenix, AZ) acts as a central supply line and is polished to create a beveled tip (Allied Multiprep Polishing Wheel, Allied High Tech Products Inc., Compton, CA). A short, additional piece of fused silica capillary, which acts as the mixing constriction or delay line, is also polished, and then glued (UV15, Master Bond Inc., Hackensack, NJ) into a larger glass capillary (320 μm ID, Polymicro Technologies, Phoenix, AZ). The supply line is secured (Microtight and Nanotight, IDEX Health and Science, Oak Harbor, WA) into custom a PEEK holder^83^, and the mixing constriction is placed over the supply line and secured. Custom laser cut centering spacers^3^ are put on all capillaries to keep everything concentric.

The Kapton observation region is added directly downstream of the mixing constriction. A small piece of glass capillary (320 μm ID, Polymicro Technologies, Phoenix, AZ) is glued (UV15, Master Bond Inc. Hackensack, NJ) onto the tip of the mixing constriction. Kapton tubing (267 μm ID, 297 μm OD, Microlumen, Oldsmar, FL) is placed inside the glass capillary and just over the tip of the mixer until it is flush with the first centering spacer. The Kapton is then secured into the glass capillary with 5-minute epoxy (ITW Devcon, Danvers, MA). The other end of the Kapton tubing is similarly glued into a piece of support glass and connected to a custom PEEK holder to complete the downstream end of the device. Both PEEK holders are mounted onto a dovetail for alignment. The Supplementary Information and Figures S2 and S3 describe the fabrication process in more detail.

### Sample delivery setup

The sample delivery setup relied on two LC-20AD Shimadzu HPLC Pumps (Shimadzu Scientific Instruments Inc., Kyoto, Japan) for robust and continuous delivery of both the crystals and the sheath. For the sample, approximately 100 m of Black PEEK Tubing (100 μm ID, Valco Instruments Company Inc., Houston, TX) was connected to the HPLC to act as a flow constrictor and build the back pressure of the flowpath to compensate for the low flow rates used (0.6-1.1 μl/min). The DJ-1 microcrystals were loaded in a 1.2 mL high pressure reservoir (Neptune Fluid Flow Systems, LLC., Knoxville, TN) and mounted on a homebuilt anti-settler with a servomotor (Digital RC Servo Motor 20KG High Torque, ANNIMOS Dsservo Technology Co. Ltd.). One end of the sample reservoir was connected directly to the supply line to reduce the dead volume of the system. Crystal flow was monitored with various cameras provided by BioCARS at the X-ray interaction region. For the sheath, about 60 m of Black PEEK Tubing was used to build back pressure, due to the slightly higher flow rates used (1-9 μl/min). Two high-pressure switching valves (Rheodyne MXP7970, IDEX Health and Science, West Henrietta, NY) were used to swap between water, buffer, and the ligand (50 mM methylglyoxal), which was loaded into a 10 mL KNAUER Variloop reservoir (KNAUER Wissenschaftliche Geräte GmbH, Berlin, Germany). The methylglyoxal was purified from pyruvaldehyde-1-dimethyl acetal as previously reported^84^ to avoid impurities present in commercial methylglyoxal. The valve system allowed for fast and efficient switching of sheath species and helped to conserve the ligand. More details are in the Supplementary Information and the entire flow path is shown in Fig. S4.

### Data collection at 14-ID-B beamline, BioCARS, APS

All time-resolved data were collected at the BioCARS 14-ID-B beamline at the APS. Two in-line undulators at this beamline with periods of 23 and 27 mm, provide high-flux polychromatic radiation at 12 keV with 5% FWHM bandwidth^63^. The mixer was mounted directly onto the goniometer, which was used as an xyz stage for alignment. The X-ray beam was focused using Kirkpatrick–Baez mirrors to beam size 22 μm vertical x 27 μm horizontal, which was a close match to the typical crystal size of 25 µm. The data were collected during the 324-bunch mode of the APS storage ring, exposing each crystal to 324 consecutive, 100 ps pulses with a total pulse train duration of 3.6 μs. The pulse trains were isolated by an ultra-fast Jülich chopper. A millisecond shutter was used to select a single opening of the chopper^63^. A Rayonix MX340-HS detector at a 9.81 Hz repetition rate was positioned 250 mm away from the sample to record diffraction images. Data were collected at 0s (no mixing) and time delays 3, 5, 10, 15, 20, and 30 sec after mixing.

### Data Processing: Precognition, Careless, CrystFEL, map generation, and difference maps

The BioCARS’ Python script^63^ was used to find hit images, defined by having at least 40-50 spots with the highest pixel count of 40-50 above the background threshold level. These selected hits were indexed using Precognition software (Renz Research, Inc.). Indexing was based on unit cell parameters and space group (P3_1_21) from an unpublished monochromatic room temperature structure of DJ-1. Based on several criteria related to the quality of geometry refinement and percentage of matched observed spots, images with multiple or mis-indexed diffraction patterns were rejected. This reduced the number of images that were processed further to approximately 50-70% of the number of images initially reported as indexed by the program. Data were integrated by Precognition to 1.7Å resolution for all time points.

After integration in Precognition, we carried out index disambiguation and merging using machine-learning tools written in Python. The source code, analysis scripts and outputs for these steps are available in a Zenodo deposition (10.5281/zenodo.10481982). We will describe the protocol briefly here. Integrated data from Space group P3_1_21 has an intrinsic a/b axis ambiguity that was resolved for each data set using a custom program (HEX-AMBI, M. Schmidt, personal communication). Another custom program, Scramble, was then used to assure consistent indexing convention for all data sets (the source code for Scramble is available on GitHub, https://github.com/Hekstra-Lab/scramble/). In this work, we used commit 746597266febb2e8dcf1a0728abe77a47e804bc8 for scramble and commit 389b3e8084cb5a443a90a3481d80d4197b5b02b3 for careless. The source code, analysis script and disambiguated output are available in our Zenodo deposition. Structure factors for each timepoint were jointly estimated using *Careless*^70^. We used a multivariate Wilson prior where each time point was statistically dependent on the initial, zero-second time point^85^. During merging, the prior correlation coefficient was set to 0.990. We used two image layers as described in the serial crystallography example^70^. Additionally, the following settings were used: Student t likelihood degrees of freedom was 32, the positional encoding frequency was 4, the mlp-wdith was 6, the test fraction was 0.05, the number of iterations was 30,000, and the half dataset repeats was 10. More details about the *Careless* settings used in this paper can be found in^86^.

In addition to Precognition-Careless processing described above, data were also processed with CrystFEL^69^. Given the wide energy bandwidth (∼5%) at BioCARS 14-ID beamline, CrystFEL-based Laue processing pipeline was used as described in Chapter 7 of Aleksandra Tolstikova’s PhD Dissertation^60^. Known room temperature unit cell parameters were used as input parameters. The same images (hits) used for indexing with Precognition were used for processing with CrystFEL. A modified CrystFEL 0.9.1 version was used for indexing with *pinkIndexer*^68^ and integration (lines 865-876 were removed from integration.c source code file as mentioned in^50^. No resolution cut off was applied at the integration step so predicted diffraction spots were integrated to the edge of the detector. The CrystFEL *indexamjig* command used for indexing and integrations is listed in Supplemental Information. Following indexing/integration, unit cell and intensity scaling steps were done (as depicted in Figure 7.12 of Tolstikova, 2020) using the measured 14-ID X-ray spectrum as input. Python scripts kindly provided by Tolstikova were used for these scaling steps. Finally, following scaling, the 0.10.2 version of CrystFEL was used to resolve indexing ambiguities with *ambigator* and to merge the data with *partialator*.

For all data, the initial DJ-1 models were refined as a rigid body against data to 2.5 Å resolution and subsequently refined to the full resolution limit using stereochemically restrained refinement of coordinates and atomic displacement parameters (ADPs) in PHENIX^87^. Riding hydrogens were added to the model, X-ray/stereochemistry and ADP weights were automatically optimized. The protein and ordered solvent models were manually optimized in COOT^88^ between PHENIX refinements. Restraints for the hemithioacetal and L-lactoylcysteine intermediates were determined using eLBOW with optimization using the eLBOW AM1 QM method^89^. Isomorphous difference (F_o_-F_o_) electron density maps were calculated between each pair of measured timepoints in PHENIX^87^ without weighting and were used in combination with polder omit mF_o_- DF_c_ electron density maps to guide the initial placement of the intermediate models and to analyze the conversion of substrate to product. Final model validation was performed in COOT^88^ and Molprobity^90^ and model statistics are in Tables S3 and S4.

## Acknowledgements

We thank Peter Zwart (Lawrence Berkeley National Laboratory) for conversations about merohedral twinning, Harrison Wang for assistance with Zenodo documentation, Graeme Winter (Diamond Light Source) for advice on DIALS processing, Aaron Brewster and Nick Sauter (Lawrence Berkeley National Laboratory) for assistance with addressing the axis ambiguity in serial crystallography data in space group P3_1_21, and Steve Meisburger (Cornell High Energy Synchrotron Source) for help with data collection and data processing.

## Data sharing

Structure factors and refined coordinates have been deposited with the Protein Data Bank (PDB) with accession codes: 9CFQ (no mixing (0s), CrystFEL-processed), 9CGA (3s mixing, CrystFEL-processed), 9CGB (5s mixing, CrystFEL-processed), 9CGD (10s mixing, CrystFEL-processed), 9CGE (15s mixing, CrystFEL-processed), 9CGF (20s mixing, CrystFEL-processed), 9CGG (30s mixing, CrystFEL-processed), 9CEI (no mixing (0s), Careless-processed), 9CFI (3s mixing, Careless-processed), 9CFM (5s mixing, Careless-processed), 9CFO (10s mixing, Careless-processed), 9CFY (15s mixing, Careless-processed), 9CFZ (20s mixing, Careless-processed), 9CG0 (30s mixing, Careless-processed), 9CMX (fixed target, no mixing), 9CMY (fixed target, methylglyoxal-mixed).

