## Supplemental material for "Resolving DJ-1 Glyoxalase Catalysis Using Mix-and-Inject Serial Crystallography at a Synchrotron"

**Table S1: CHESS Data Collection Statistics**

| | | |
|---|---|---|
| **X-ray Source** | Cornell High Energy Synchrotron Source, ID-7-B-2 | |
| **X-ray energy (keV)** | 12.8 (0.6% bandwidth) | |
| **Transmission** | 52% | |
| **Temperature (K)** | 293 | |
| **Detector** | Eiger2 X 16M | |
| **Concentration of methylglyoxal for soaking** | 0 mM | 40 mM |
| **Oscillation range** | 0-0.25° | |
| **Number of frames per crystal** | 5 | |
| **Total exposure time per crystal** | 50 ms | |
| **Sample** | No mixing | Methylglyoxal |
| **Total number of oscillation series collected** | 680 | 632 |
| **Number of hits** | 367 | 414 |
| **Hit rate** | 54.0% | 65.5% |
| **Space group** | $P3_121$ | |
| **a, b, c (Å)** | 76.23, 76.23, 76.39 | |
| **α, β, γ (°)** | 90, 90, 120 | |
| **Number indexed** | 197 | 228 |
| **Indexing rate** | 53.7% | 55.1% |
| **Number integrated** | 165 | 196 |
| **Number merged** | 157 | 187 |
| **Resolution (Å)** | 66.10-1.63 (1.66-1.63) | 66.02-1.69 (1.72-1.69) |
| **$\langle I/\sigma(I)\rangle$** | 8.5 (0.7) | 6.6 (0.7) |
| **$R_{merge}$** | 0.224 (4.819) | 0.295 (5.560) |
| **Number of integrated intensities (observations)** | 366392 | 371031 |
| **Number of unique reflections** | 32118 (1632) | 28986 (1428) |
| **Redundancy** | 11.4 (11.5) | 12.8 (12.9) |

| | | |
|---|---|---|
| **Completeness (%)** | 98.61 (99.69) | 99.31 (99.86) |
| **$CC_{1/2}$** | 0.990 (0.260) | 0.976 (0.066) |

**Table S2: MISC Data Collection and Processing Statistics**

| **X-ray source** | Advanced Photon Source, 14-ID-B beamline | | | | | | |
|---|---|---|---|---|---|---|---|
| **Sample** | DJ-1 with methylglyoxal mixing | | | | | | |
| **Energy range** | 10-12.2 keV | | | | | | |
| **Temperature (K)** | 293 | | | | | | |
| **Detector** | Rayonix MX340-HS | | | | | | |
| **Space Group** | $P3_121$ | | | | | | |
| **a, b, c (Å)** | 76.03, 76.03, 76.14 | | | | | | |
| **α, β, γ (°)** | 90, 90, 120 | | | | | | |
| ***All Datasets: Hit Finding by pyPrecognition*** | | | | | | | |
| **Timepoint** | **0 s** | **3 s** | **5 s** | **10 s** | **15 s** | **20 s** | **30 s** |
| **No. of frames** | 89,887 | 25,000 | 25,000 | 20,000 | 30,000 | 23,000 | 20,000 |
| **No. of hits** | 5,332 | 1,695 | 1,169 | 1,518 | 1,864 | 998 | 1,051 |
| **Hit rate** | 5.93% | 6.78% | 4.68% | 7.59% | 6.21% | 4.34% | 5.26% |
| ***Precognition Alone*** | | | | | | | |
| **Indexed images** | 1748 | 512 | 409 | 653 | 1050 | 375 | 354 |
| **Indexing rate** | 32.78% | 30.21% | 34.99% | 43.02% | 56.33% | 37.58% | 33.68% |
| **Integrated images** | 507 | 279 | 235 | 306 | 505 | 259 | 211 |
| **Merged images** | 506 | 278 | 234 | 306 | 505 | 253 | 211 |
| **Resolution cutoff for integration (Å)** | 1.7 | 1.7 | 1.7 | 1.7 | 1.7 | 1.7 | 1.7 |
| ***Precognition + Careless*** | | | | | | | |
| **Merged images (from Precognition)** | 506 | 278 | 234 | 306 | 505 | 253 | 211 |
| **Resolution range (Å)*** | 65.84-1.77 (1.83-1.77) | 65.84-1.77 (1.83-1.77) | 65.84-1.77 (1.83-1.77) | 65.84-1.77 (1.83-1.77) | 65.84-1.77 (1.83-1.77) | 65.84-1.77 (1.83-1.77) | 65.84-1.77 (1.83-1.77) |
| **<I/σ(I)>** | 17.21 (2.22) | 13.33 (1.69) | 12.67 (1.56) | 15.26 (1.80) | 20.43 (2.39) | 10.93 (1.49) | 12.35 (1.53) |

| | | | | | | | |
|---|---|---|---|---|---|---|---|
| **Integrated intensities** | 1,642,115 | 901,776 | 759,698 | 992,682 | 1,639,462 | 824,105 | 685,193 |
| **Unique reflections** | 25,220 (2,459) | 25,216 (2,459) | 25,208 (2,455) | 25,222 (2,457) | 25,225 (2,460) | 25,219 (2,458) | 25,209 (2,448) |
| **Redundancy** | 65.1 (21.9) | 35.8 (12.0) | 30.1 (10.2) | 39.4 (13.4) | 65.0 (22.0) | 32.7 (11.0) | 27.2 (9.2) |
| **Completeness (%)** | 99.85 (99.84) | 99.83 (99.84) | 99.78 (99.68) | 99.83 (99.76) | 99.86 (99.88) | 99.83 (99.80) | 99.79 (99.47) |
| **$CC_{1/2}$** | 0.98 (0.33) | 0.98 (0.35) | 0.98 (0.34) | 0.99 (0.34) | 0.99 (0.35) | 0.98 (0.35) | 0.98 (0.34) |
| ***CrystFEL*** | | | | | | | |
| **Number of hits (from pyPrecognition )** | 5,332 | 1,695 | 1,169 | 1,518 | 1,864 | 998 | 1,051 |
| **Indexed & merged images** | 5,073 | 1,561 | 1,113 | 1,477 | 1,824 | 949 | 1,007 |
| **Indexing and merging rate** | 95.14% | 92.09% | 95.21% | 97.30% | 97.85% | 95.09% | 95.81% |
| **Resolution range (Å)** | 32.96-1.77 (1.80-1.77) | 32.96-1.77 (1.80-1.77) | 32.96-1.77 (1.80-1.77) | 32.96-1.77 (1.80-1.77) | 32.96-1.77 (1.80-1.77) | 32.96-1.77 (1.80-1.77) | 32.96-1.77 (1.80-1.77) |
| **$R_{split}$ (%)**** | 5.98 | 11.21 | 11.67 | 10.10 | 9.96 | 15.03 | 13.75 |
| **$\langle I/\sigma(I)\rangle$** | 7.74 (0.43) | 4.22 (0.23) | 3.98 (0.23) | 4.68 (0.31) | 5.24 (0.37) | 3.03 (0.15) | 3.64 (0.27) |
| **Integrated intensities (observations)** | 6,151,235 | 2,393,539 | 1,650,294 | 2,382,889 | 2,963,509 | 1,443,367 | 1,515,860 |
| **Unique reflections** | 25,005 (2426) | 24,997 (2428) | 24,994 (2427) | 24,998 (2428) | 25,016 (2431) | 24,990 (2428) | 24,993 (2428) |
| **Redundancy** | 246.00 (206.6) | 96.75 (91.5) | 66.03 (63.3) | 95.32 (90.5) | 118.46 (112.8) | 57.76 (55.0) | 60.65 (58.4) |
| **Completeness (%)** | 99.03 (98.62) | 99.01 (98.70) | 98.99 (98.66) | 99.01 (98.70) | 99.08 (98.82) | 98.98 (98.70) | 98.98 (98.70) |
| **$CC_{1/2}$** | 0.99 (0.05) | 0.98 (-0.03) | 0.98 (0.03) | 0.99 (0.05) | 0.98 (0.09) | 0.98 (0.04) | 0.98 (0.04) |

*An input parameter to Careless, determined based on keeping $CC_{1/2}$ above 0.3 at highest resolution. All Careless stats are based on this cutoff

All CrystFEL statistics provided here, from $R_{split}$ to $CC_{1/2}$, were calculated to 1.77 Å to facilitate comparisons with Careless statistics. Values in parenthesis are for the highest resolution shell. Different cutoffs were used for refinement, as detailed in Tables S4 and S5.

**Table S3: Model Statistics, CHESS fixed target datasets**

| **Sample** | No substrate | 40 mM methylglyoxal |
|---|---|---|
| | 9CMX | 9CMY |
| **Refinement Program** | PHENIX 1.19.2-4158 | |
| **Resolution range** | 38.16 - 1.63 (1.69 - 1.63) | 38.12 - 1.69 (1.75 - 1.69) |
| **Completeness (%)** | 98.15 (97.52) | 99.04 (98.04) |
| **Working set (no. reflections)** | 31968 (3150) | 28978 (2807) |
| **Test set (no. reflections)** | 2005 (194) | 1931 (199) |
| **$R_{work}$** | 0.1437 (0.3033) | 0.1572 (0.2940) |
| **$R_{free}$** | 0.1672 (0.3425) | 0.1751 (0.3342) |
| **Number of non-hydrogen atoms** | 1574 | 1566 |
| **Protein** | 1472 | 1474 |
| **Ligands** | 0 | 0 |
| **Water** | 102 | 92 |
| **Protein residues** | 188 | 188 |
| ***Average RMS deviation*** | | |
| **Bonds (Å)** | 0.011 | 0.007 |
| **Angles (°)** | 1.14 | 0.78 |
| ***Average B-factors ($Å^2$)*** | | |
| **Protein** | 30.48 | 32.06 |
| **Ligands** | - | - |
| **Water** | 43.06 | 40.79 |
| **Clashscore** | 0.99 | 1.66 |
| ***Ramachandran plot*** | | |
| **Favored (%)** | 99.45 | 98.92 |
| **Allowed (%)** | 0.55 | 0.54 |
| **Outliers (%)** | 0.00 | 0.54 |

**Table S4: Model Statistics APS Laue MISC datasets, CrystFEL processing**

| Sample | CrystFEL_0s | CrystFEL_3s_HTA | CrystFEL_5s_HTA | CrystFEL_10s_LC | CrystFEL_15s_LC | CrystFEL_20s_HTA | CrystFEL_30s_LC |
|---|---|---|---|---|---|---|---|
| **PDB Code** | 9CFQ | 9CGA | 9CGB | 9CGD | 9CGE | 9CGF | 9CGG |
| **Refinement Program** | PHENIX 1.19.2-4158 | | | | | | |
| **Resolution range (for refinement)** | 32.96-1.90 (1.97-1.90) | 32.96-2.01 (2.08-2.01) | 32.96-2.01 (2.08-2.01) | 32.96-1.97 (2.04-1.97) | 32.96-1.90 (1.97-1.90) | 32.96-2.06 (2.13-2.06) | 32.96-2.01 (2.08-2.01) |
| **Completeness (%)** | 99.09 (99.19) | 99.05 (99.05) | 99.04 (99.11) | 99.03 (98.88) | 99.11 (99.24) | 98.98 (98.91) | 99.00 (98.99) |
| **Working set (no. reflections)** | 20292 (1964) | 17202 (1667) | 17200 (1668) | 18236 (1763) | 20296 (1965) | 15979 (1549) | 17194 (1666) |
| **Test set (no. reflections)** | 1017 (113) | 861 (96) | 861 (96) | 902 (81) | 1016 (113) | 797 (86) | 861 (96) |
| **$R_{work}$** | 0.1497 (0.3080) | 0.1577 (0.3070) | 0.1623 (0.3019) | 0.1490 (0.2824) | 0.1532 (0.3231) | 0.1726 (0.3074) | 0.1630 (0.2976) |
| **$R_{free}$** | 0.1661 (0.3166) | 0.1790 (0.3027) | 0.1877 (0.3276) | 0.1768 (0.2960) | 0.1760 (0.3636) | 0.1890 (0.3109) | 0.1822 (0.3121) |
| **Number of non-hydrogen atoms** | 1578 | 1553 | 1582 | 1566 | 1557 | 1562 | 1549 |
| **Protein** | 1442 | 1442 | 1450 | 1437 | 1435 | 1435 | 1431 |
| **Ligands** | 0 | 5 | 5 | 5 | 5 | 5 | 5 |
| **Water** | 136 | 106 | 127 | 124 | 117 | 122 | 113 |
| **Protein residues** | 188 | 188 | 188 | 188 | 188 | 188 | 188 |
| **Average RMS deviation** | | | | | | | |
| **bonds (Å)** | 0.006 | 0.003 | 0.002 | 0.006 | 0.003 | 0.002 | 0.002 |
| **angles (°)** | 0.61 | 0.62 | 0.53 | 0.73 | 0.62 | 0.51 | 0.51 |
| **Average B-factors ($Å^2$)** | | | | | | | |
| **Protein** | 31.56 | 34.13 | 33.47 | 33.23 | 33.37 | 34.02 | 33.23 |
| **Ligands** | - | 39.88 | 30.90 | 37.18 | 37.85 | 33.67 | 33.96 |
| **Water** | 45.34 | 44.52 | 45.87 | 48.59 | 45.64 | 45.58 | 46.23 |
| **Clashscore** | 2.37 | 1.69 | 0.67 | 1.36 | 1.02 | 1.02 | 1.36 |
| **Ramachandran plot** | | | | | | | |
| **Favored (%)** | 99.45 | 98.92 | 98.92 | 98.92 | 98.92 | 98.92 | 98.92 |
| **Allowed (%)** | 0.55 | 0.54 | 0.54 | 0.54 | 0.54 | 0.54 | 0.54 |
| **Outliers (%)** | 0 | 0.54 | 0.54 | 0.54 | 0.54 | 0.54 | 0.54 |

**Table S5: Model Statistics APS Laue MISC datasets, Careless processing**

| ***APS Laue MISC datasets, Careless processing*** | | | | | | | |
|---|---|---|---|---|---|---|---|
| **Sample** | Careless_0s | Careless_3s _HTA | Careless_5s _HTA | Careless_10s _LC | Careless_15s _LC | Careless_20s _HTA | Careless_30s _LC |
| **PDB code** | 9CEI | 9CFI | 9CFM | 9CFO | 9CFY | 9CFZ | 9CG0 |
| **Refinement Program** | PHENIX 1.19.2-4158 | | | | | | |
| **Resolution range** | 38.01-1.77 (1.83-1.77) | 32.96-1.77 (1.83-1.77) | 32.96-1.77 (1.83-1.77) | 49.80-1.77 (1.83-1.77) | 49.80-1.77 (1.83-1.77) | 38.01-1.77 (1.83-1.77) | 38.01-1.77 (1.83-1.77) |
| **Completeness (%)** | 99.86 (99.84) | 99.83 (99.84) | 99.80 (99.68) | 99.85 (99.76) | 99.86 (99.88) | 99.84 (99.80) | 99.80 (99.47) |
| **Working set (no. reflections)** | 25190 (2459) | 25183 (2459) | 25176 (2455) | 25188 (2457) | 25191 (2460) | 25185 (2458) | 25174 (2448) |
| **Test set (no. reflections)** | 1145 (155) | 1145 (155) | 1145 (155) | 1145 (155) | 1145 (155) | 1145 (155) | 1143 (154) |
| **$R_{work}$** | 0.2044 (0.3299) | 0.2045 (0.3264) | 0.2046 (0.3256) | 0.2004 (0.3273) | 0.2008 (0.3295) | 0.2036 (0.3269) | 0.2031 (0.3276) |
| **$R_{free}$** | 0.2350 (0.3357) | 0.2347 (0.3336) | 0.2398 (0.3567) | 0.2312 (0.3456) | 0.2296 (0.3491) | 0.2369 (0.3391) | 0.2361 (0.3459) |
| **Number of non-hydrogen atoms** | 1578 | 1554 | 1582 | 1566 | 1557 | 1562 | 1549 |
| **Protein** | 1442 | 1442 | 1450 | 1437 | 1435 | 1435 | 1431 |
| **Ligands** | 0 | 5 | 5 | 5 | 5 | 5 | 5 |
| **Water** | 136 | 107 | 127 | 124 | 117 | 122 | 113 |
| **Protein residues** | 188 | 188 | 188 | 188 | 188 | 188 | 188 |
| ***Average RMS deviation*** | | | | | | | |
| **Bonds (Å)** | 0.005 | 0.017 | 0.019 | 0.016 | 0.009 | 0.017 | 0.009 |
| **Angles (°)** | 0.734 | 1.292 | 1.774 | 1.502 | 0.933 | 1.524 | 1.107 |
| ***Average B-factors (Å²)*** | | | | | | | |
| **Protein** | 16.94 | 14.77 | 16.22 | 14.34 | 14.84 | 14.55 | 15.07 |
| **Ligands** | - | 18.82 | 26.14 | 17.07 | 18.89 | 23.79 | 19.28 |
| **Water** | 31.00 | 25.16 | 31.39 | 26.75 | 27.14 | 27.57 | 28.21 |
| **Clashscore** | 1.69 | 3.38 | 3.36 | 2.37 | 1.02 | 1.70 | 1.36 |
| ***Ramachandran plot*** | | | | | | | |
| **Favored (%)** | 98.91 | 98.92 | 98.39 | 98.39 | 98.39 | 97.85 | 98.39 |
| **Allowed (%)** | 1.09 | 0.54 | 1.08 | 1.08 | 1.08 | 1.61 | 1.08 |
| **Outliers (%)** | 0.00 | 0.54 | 0.54 | 0.54 | 0.54 | 0.54 | 0.54 |

Values in parentheses are for the highest resolution shell

**Supplemental Methods and Figures**

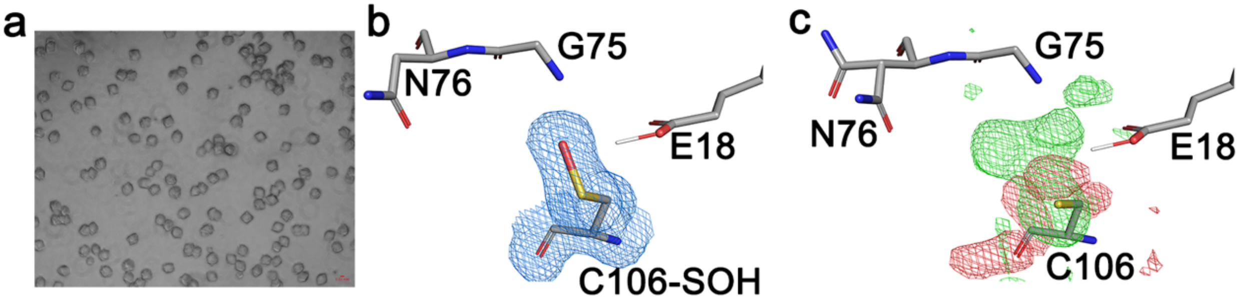


**Figure S1. Fixed target serial crystallography shows sulfenic acid formation and evidence of methylglyoxal modification at C106.** Panel (a) shows ~25 $\mu$m DJ-1 microcrystals grown using seeding with scale bar in red at lower right. Panel (b) shows a $2mF_o$-$DF_c$ electron density map contoured at $1.0\sigma$ (blue) with a cysteine sulfenic acid (Cys-SOH) modeled at Cys106, the active site nucleophile. The persistence of this modification, even in the presence of DTT reductant, suggests that it is due to X-ray photooxidation. Panel (c) shows a $F_o$(methylglyoxal)-$F_o$(free) difference electron density map contoured at $+3\sigma$ (green) and $-3\sigma$ (red) around Cys106, indicating modification by methylglyoxal. Although the nature of the experiment does not allow detailed timescale information to be determined, the formation of a putative intermediate upon mixing with methylglyoxal substrate suggests catalytic activity *in crystallo*. In addition, the ability of Cys106 to be modified by methylglyoxal prior to exposure to the X-ray beam, which requires a reduced Cys106, supports photooxidation as the explanation for Cys106-SOH formation in the free enzyme (b).

*Mixer and Flow Cell Fabrication*

The microfluidic mixer is fabricated as described in our previous work (Plumridge *et al.*, 2018; Calvey *et al.*, 2019). Briefly, a piece of capillary that acts as the mixing and delay line is polished to a tip and glued inside a larger piece of glass at the upstream end. Kapton centering spacers (Calvey *et al.*, 2016) are placed on the capillary to keep everything concentric. The supply line for the crystals (0.5-1 meter length of capillary) is also polished to a tip with spacers and secured into the upstream custom PEEK holder with a coned fitting. Next, the larger piece of glass with the mixing line is installed over the supply line and secured to the PEEK holder with another fitting (Figure S2a). The spacing between the end of the supply line and the start of the mixing line is adjusted to be ~100 μm. The assembly is then placed on a dovetail rail.

The coupling of the mixer to the Kapton tubing to create the flow cell is a new design for this device. Since the Kapton tubing is quite flexible, a piece of support glass (ID 320 μm) is used to stabilize the connection at both the upstream and downstream ends. First, the downstream support glass is secured with a fitting to the downstream custom PEEK holder. Next, the Kapton tubing is placed inside the support glass and the assembly is placed on the dovetail rail (Figure S2a). The upstream support glass is placed over the entering spacers on the downstream end of the mixing capillary (Figure S2b). Then, the Kapton tubing is slid inside the upstream support glass and over the tip of the capillary until it is touching the first spacer by moving both the downstream PEEK holder and the Kapton tubing itself (Figure S2c). The position of the upstream support glass is adjusted to minimize the overlap between the glass and the Kapton. Lastly, the support glass is secured onto the mixing line using UV curable epoxy (UV-15) and the Kapton tubing is bonded to both support glasses using 5-minute Epoxy as Kapton is not UV transparent (Figure S2d). A small needle is used to apply the epoxy to minimize the size of the glue blob, especially on the upstream side.

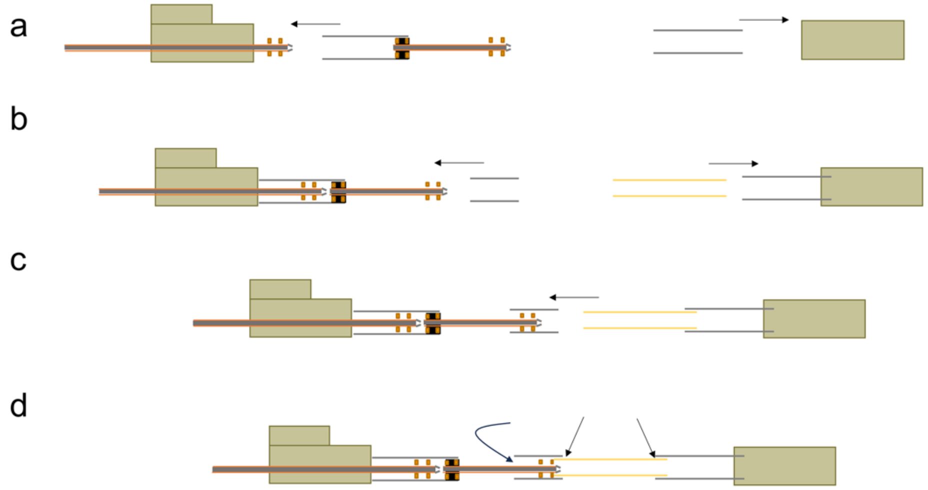


**Figure S2: Schematic of the mixer and flow cell fabrication**. Panel a: On the upstream side, place the mixer body (glass and constriction) over the supply line. On the downstream side, secure the support glass into the PEEK body. Panel b: On the upstream side, place support glass over the end of the constriction. On the downstream side, slide the Kapton tubing into the support glass. Panel c: Slide the Kapton tubing into the upstream support glass. Panel d: Glue all junctions (positions of arrows)

*Timepoint Calculation*

The final timepoint measured during data collection is calculated as described in[1,2]. After the sample stream is flow focused down to a thin stream, the ligand rapidly diffuses into the sample stream and into the crystal to initiate the reaction. This increases the concentration of the ligand in the sample stream, but with a radial spread because it takes longer for the ligand to reach the center of the sample stream. The system is defined as fully mixed when the concentration of ligand and the concentration of protein inside the crystal are equimolar at the center of the sample stream. At this point, the system is given an average age and an uncertainty based on how rapidly the ligand diffuses into the sample stream. Importantly for this work, the width of the sample stream was required to be at least 20-25 μm to match the size of the crystal, and this places a lower limit on fast mixing.

Once the system is fully mixed, it continues to age for a set additional delay. The first additional delay is accrued inside the mixing capillary, simply based on the flow speed and the remaining length of the mixer. The spread in speed due to the parabolic flow profile is taken into account as an additional contributor to the uncertainty. It is also important to note that the ligand continues to diffuse into the sample stream, eventually causing the ligand to be in excess of the protein concentration. Next, the freshly mixed species enters the Kapton flow cell, and additional delay times can be gained based on where the X-ray beam is positioned relative to the tip of the mixing line. This distance, which can vary from ~0.3 mm - 8 mm (limit of the motor at BioCARS) and allows for considerable flexibility. The additional delay time accrued in the Kapton tubing is similarly calculated based on the flow speed and the distance traveled, while still accounting for the spread in speed for the uncertainty.

In short, the final timepoint is a sum of the following: the average age of the sample when fully mixed, the delay inside the mixing line, and the delay inside the Kapton tubing with each contributing to the total uncertainty of the timepoint. The ligand flowrate, sample flowrate, and position of the X-ray beam can all be varied to reach different timepoints within a single flow cell, allowing access to a wide range of timepoints in a single device.

*Expansion Region Mixer*

In addition to the design described above, we have a second design, the Expansion Region Mixer, to facilitate data acquisition at long timepoints without making the devices unreasonably long and more fragile. The schematic of the Expansion Region Mixer is shown in Figure S3. Instead of having a single mixing and delay line (Figure S3a), the Expansion Region Mixer (Figure S3b) consists of three parts: a short mixing line (100-120 μm ID), an expansion region delay section (320 μm ID), and a final delay and focusing line (100-120 μm ID). The fabrication of the expansion region mixer follows the same steps as above and is depicted in Figure S3. The only extra steps involve gluing the second delay line to the expansion region and then gluing the expansion region onto the first delay line. Spacers are used at each of these junctions to keep everything concentric.

In the first mixing line, everything proceeds as described above, and the system becomes fully mixed at some point in this section. Next, the sample enters the expansion region, and due to the larger inner diameter, the flow speed decreases. This slow down allows the system to age more efficiently as achieving the same delay time in a standard capillary would require a longer distance. It is essential for the system to be fully mixed before entering the expansion region as the increase in the width of the sample stream also results in a slowdown in diffusion of the ligand into the sample stream. After the expansion region, the sample enters a second delay line, which allows for additional flexibility in reaching the overall timepoint and also focuses the crystal stream back down before it enters the Kapton region. An additional delay is still typically accrued in the Kapton region, but generally measurements are done closer to the mixer tip as the sample is already aged more.

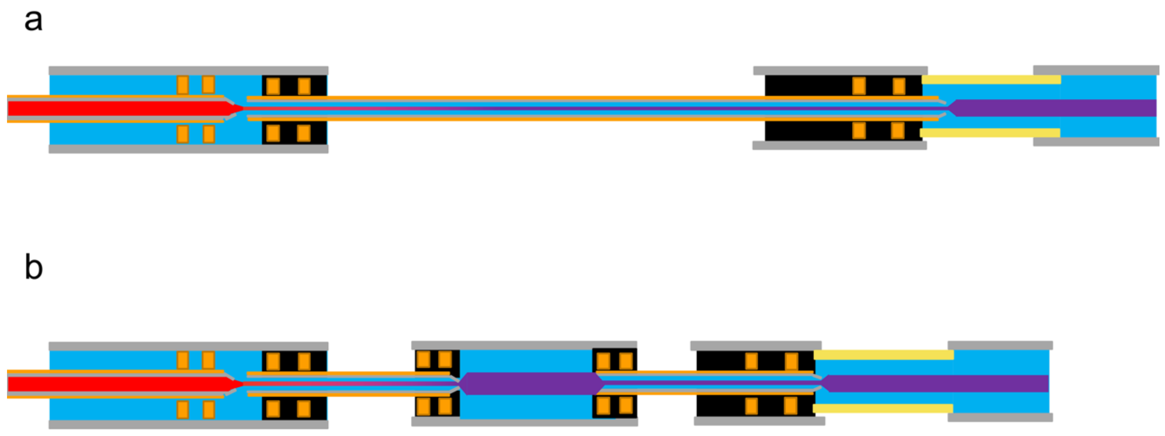


**Figure S3: Schematic of standard and expansion region microfluidic mixers.** A. Standard microfluidic mixer with a long constriction coupled to the Kapton observation region (same as depicted in Fig. 2a). B. Expansion region microfluidic mixer. A standard constriction is coupled to a larger glass section to reduce the flow speed and accrue a greater delay in a shorter distance. Next, a second constriction is added to better focus the crystals before entering the Kapton observation region. The overall footprint of the device is shorter than the standard microfluidic mixer that could access that time range and extends the timepoints achievable in these devices.

Briefly, the final timepoint for the expansion region device has four contributions: the average age of the sample when fully mixed, the delay from the expansion region, the delay from the extra delay and focusing line, and the delay inside the Kapton tubing with each contributing to the total uncertainty of the timepoint. The benefit of the expansion region mixer is that longer timepoints can be reached in a shorter distance, which can help prevent the device from being unreasonably long and more fragile. To reach the same timepoint in the expansion region and standard mixer, the flowrates are slightly higher for the expansion region mixer. Although in some cases this may be prohibitive, it generally helps with experimental efficiency because higher sample flowrates correlate with higher hit rates and faster data collection.

**Table S6**: Mixer and Timepoint Details

| 80 mm, 100 µm ID Mixer | | | | |
|---|---|---|---|---|
| Timepoint (s ± uncertainty) | Sample Flow Rate (µL/min) | Sheath Flow Rate (µL/min) | X-ray Beam Position (mm from tip) | Final Concentration of methylglyoxal (mM) |
| 3 ± 0.072 | 1.1 | 6.7 | 2.3 | 42 |
| 5 ± 0.116 | 0.8 | 4 | 2.8 | 41 |
| 20 ± 1.504 | 0.6 | 1 | 6.7 | 32 |
| Expansion Region Mixer (35 mm, 100 µm ID; 17 mm, 320 µm ID; 20 mm, 100 µm ID) | | | | |
| Timepoint (s ± uncertainty) | Sample Flow Rate (µL/min) | Sheath Flow Rate (µL/min) | X-ray Beam Position (mm from tip) | Final Concentration of methylglyoxal (mM) |
| 5 ± 0.087 | 1 | 9 | 0.45 | 45 |
| 10 ± 0.228 | 0.8 | 4.9 | 0.56 | 42 |
| 30 ± 2.056 | 0.6 | 1.2 | 6.5 | 34 |

*Final Mixer Designs*

Both mixers cover similar time ranges, but the flow rates needed vary due to the difference in the overall mixing and delay times. The 80 mm mixer uses overall lower flow rates because its delay region is shorter, but it is challenging to go beyond 20 s, whereas the expansion region mixer uses slightly higher flow rates and can accommodate longer timepoints. Thus, there is a lot of flexibility in the device design stage to balance sample/ligand consumption, timepoint uncertainty, and accessible time ranges.

*Mixer Operation: Fluidics*

The mixer operation is quite straightforward and the fluidic consist of two components, one for the sample and one for the sheath as depicted in Figure S4. Both utilize Shimadzu HPLC pumps to drive the flow, and the first component of both sides is 60-100 m of 100µm PEEK tubing to build the back pressure of the system and accommodate the low flowrates. From flow tests using Elveflow flow meters to monitor the stability of the flow, a minimum back pressure of ~100 PSI is sufficient. For the sample side, the HPLC is either connected directly to the supply line or to a sample reservoir. When the HPLC is connected directly to the supply line, water flows through, which is important at startup or for rinsing the line. To flow crystals into the line, the HPLC is instead connected to the back end of a 1.2 mL reservoir. The front end of the reservoir is loaded with crystals, and when the back end is connected to the HPLC, water pushes against a piston to push the crystals out of the reservoir and into the supply line. Importantly, the reservoir is connected directly to the supply line to avoid having the crystals travel through any valves, minimizing the chance of flow issues. Additionally, the reservoir is mounted on an anti-settler device to keep rotating the reservoir to prevent crystal settling inside the reservoir. Switching between water and crystals is done manually by swapping out the sample reservoir for a standard fluidic union.

The sheath side fluidics utilize two selection valves in addition to the HPLC and the back pressure lines. The selection valves allow for automatic switching between flowing water or the ligand to the sample cell. The HPLC line connects to the center of one of the selection valves and then, depending on which port the valve is set to, the water flows directly to the second selection valve and into the sample cell or the water flows to a sheath reservoir. The sheath reservoir has a 10 mL volume, and, just like the sample reservoir, the water from the HPLC flows into the backside of the reservoir to push a piston that drives the ligand out of the reservoir and into the next line. The line is connected to the second selection valve, and when the correct port is set, it will flow to the sample cell.

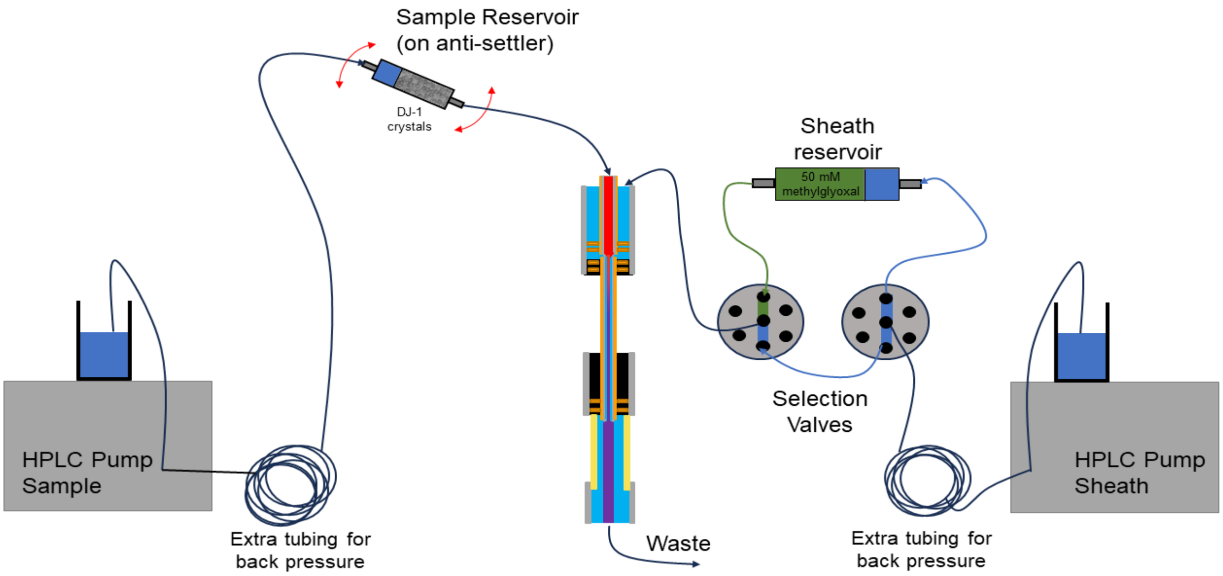


**Figure S4: Schematic of the fluidic setup to drive the sample and sheath**. For the sample, the HPLC is connected to extra tubing to increase the back pressure of the system. Then the sample reservoir, on an anti-settler, is directly connected to the supply line of the flow cell. For the sheath, the HPLC is again connected to extra tubing to help with the back pressure. Next, two selection valves choose between flowing water or ligand to the sheath line of the flow cell.

Mixer operation for data collection proceeds as followed:

1. Flow water through all sample and sheath lines, including both sides of the selection valve for at least 5 minutes
2. Position the sample cell so the X-ray beam is at the correct distance from the tip for data collection
3. Load ligand into the sheath reservoir
4. Connect the sheath reservoir to the selector valves and set the flowrate to 5-10 μL/min and start keeping track of how much volume is flown
5. Switch the selector valves to the sheath reservoir position
6. Flush the lines and sample cell with ligand for 5-10 minutes
7. Filter fresh crystals and load into the sample reservoir
8. Mount sample reservoir on anti-settler and connect lines to the reservoir and set the flowrate to 1-3 μL/min and start keeping track of how much volume is flown
9. Turn anti-settler on
10. Wait a few minutes for the crystals to get through the lines and use cameras to see crystals appear
11. Adjust flowrates to the ones required for the timepoint of interest
12. Close hutch
13. Start data collection

To switch to a new timepoint:

1. Confirm that both the sheath reservoir and sample reservoir are full enough for a subsequent measurement. If not, see the instructions below to end data collection, and then follow the procedure above for refilling reservoirs and re-starting data collection.
2. Adjust the sheath flowrate
3. Adjust the sample flowrate
4. Move the sample cell to the correct position for the desired timepoint
5. Start data collection

To end data collection:

1. Open the hutch
2. Set the sheath flow up to 5-10 μL/min
3. Swap the crystal reservoir for a fluidic union and set the flow to 1-3 μL/min
4. Rinse the crystal line for 5-10 min
5. Swap the selector valve to water
6. Rinse the sheath line for 5-10 min
7. Replace the sheath reservoir with a fluidic union and swap the selector valve back to that port
8. Rinse the selector valve ports and lines with water
9. Empty both the sample and sheath reservoirs
10. Clean both with 2% Hellmanex six times, and with water three times

***Data Analysis: CrystFEL***

Indexamajig command for CrystFEL indexing and integration:

indexamajig -i files.lst -o 0-0s.stream -g mccd.geom -p DJ1.cell --peaks=peakfinder9 --threshold=5 --min-snr=4.5 --local-bg-radius=3 --min-snr-biggest-pix=2 --min-snr-peak-pix=1 --min-peaks=20 --indexing=pinkIndexer-latt --pinkIndexer-considered-peaks-count=4 --pinkIndexer-angle-resolution=4 --pinkIndexer-max-resolution-for-indexing=0.45 --pinkIndexer-refinement-type=5 --pinkIndexer-tolerance=0.03 --no-retry --no-refine --no-check-peaks --no-check-cell --fix-profile-radius=0.001e8 --int-radius=4,6,8 --no-non-hits-in-stream --pinkIndexer-thread-count=20 -j 6

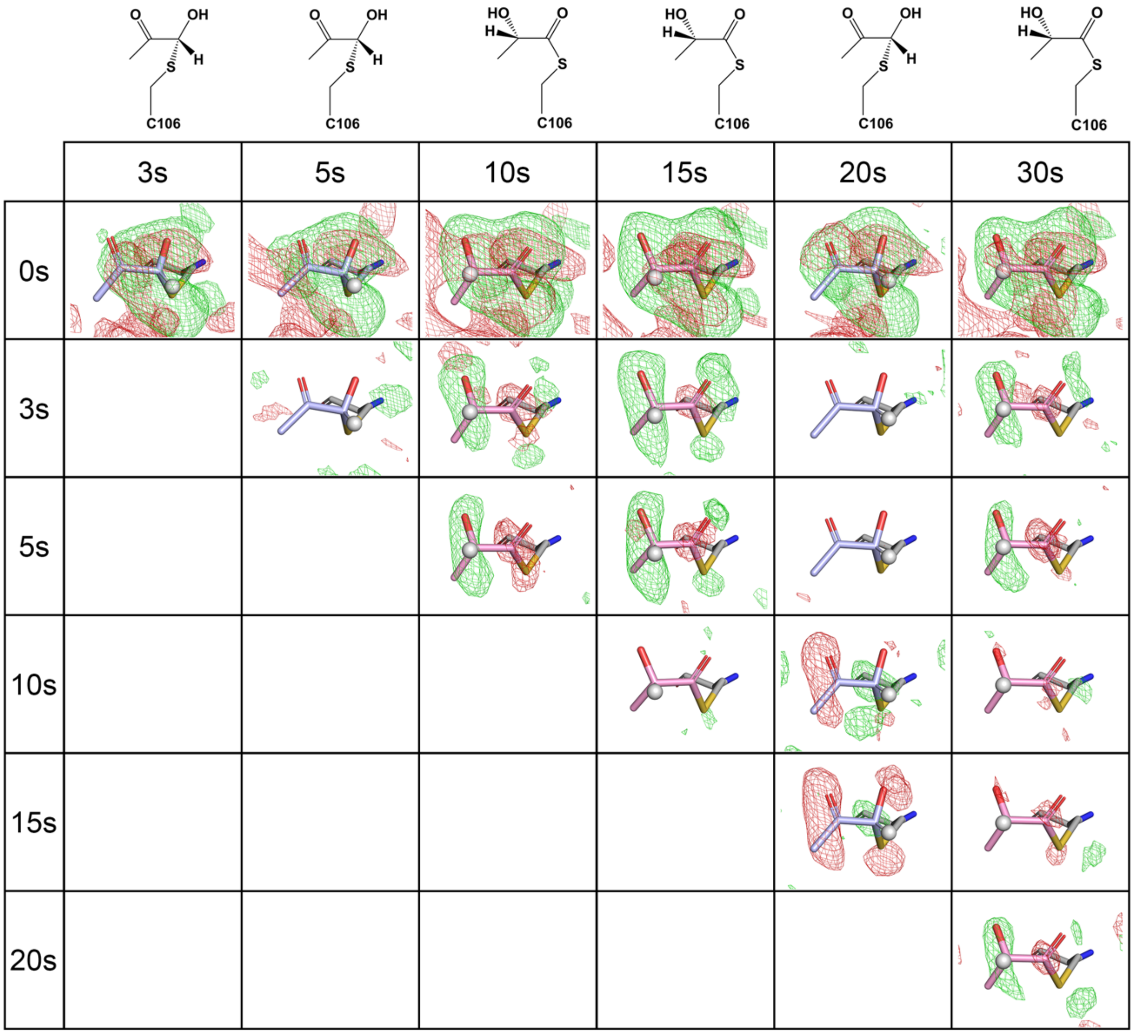


**Figure S5: Matrix of $F_o$-$F_o$ electron density maps in DJ-1 active site, Careless-processed data.** $F_o$-$F_o$ electron density maps are contoured at +3σ (green) and -3σ (red) around the intermediate at Cys106. Each position in the matrix shows the map calculated by subtracting the shorter from the longer timepoint. The chemical structures for the intermediates modeled at each timepoint are shown in the top row. The featureless maps for the $F_o$(5s)-$F_o$(3s), $F_o$(15s)-$F_o$(10s), $F_o$(20s)-$F_o$(3s), $F_o$(20s)-$F_o$(5s), $F_o$(30s)-$F_o$(10s), and $F_o$(30s)-$F_o$(15s) pairs indicate that the species present at these timepoints are identical. Therefore, we propose that DJ-1 catalyzes nearly two turnovers in the crystal, with the hemithioacetal present at 3,5, and 20s and the L-lactoylcysteine present at 10,15, and 30s. The matrix is symmetric about the diagonal, so only one unique half is shown.

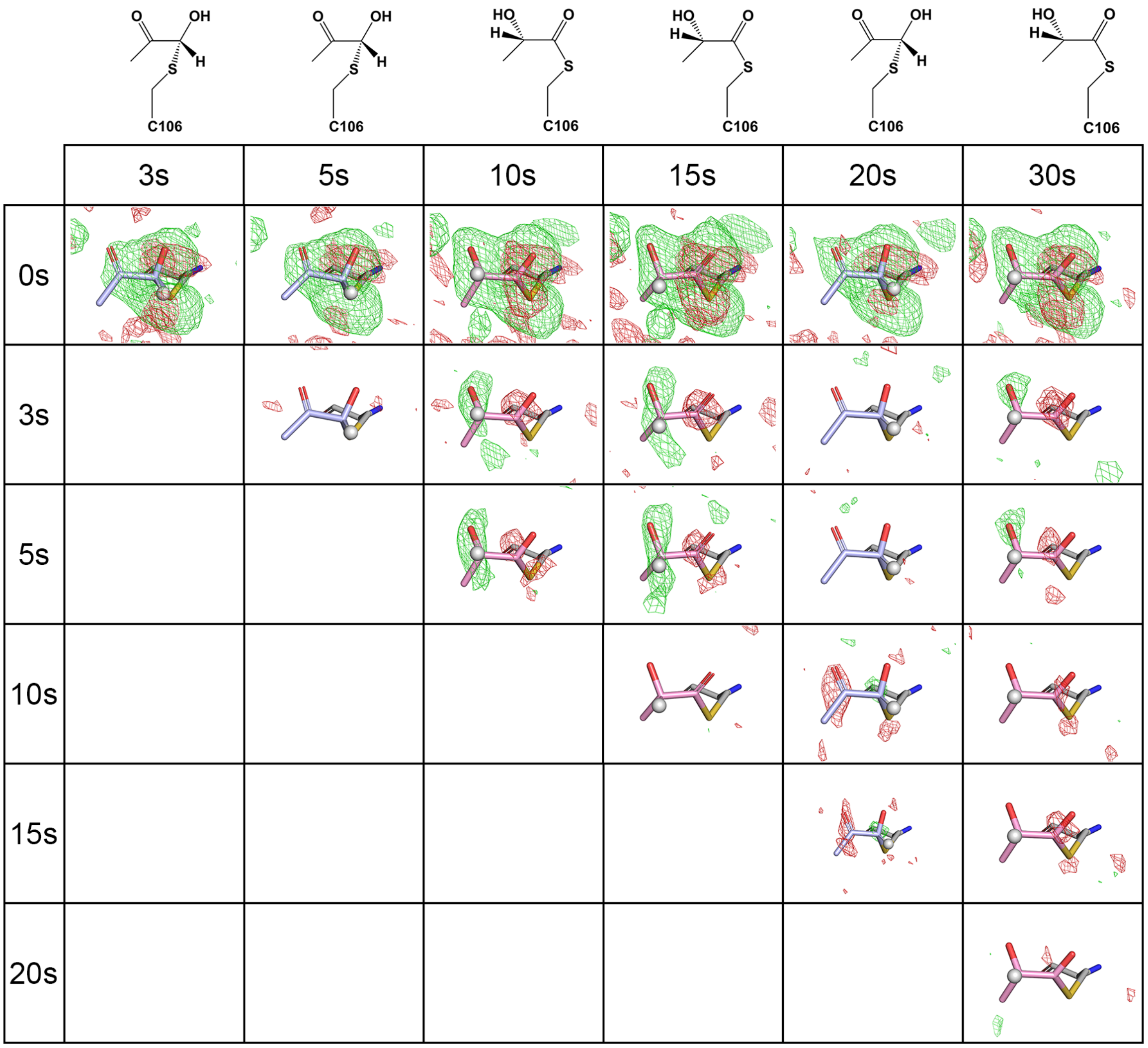


**Figure S6: Matrix of $F_o$-$F_o$ electron density maps in DJ-1 active site, CrystFEL-processed data.** $F_o$-$F_o$ electron density maps are contoured at +3σ (green) and -3σ (red) around the intermediate at Cys106. Each position in the matrix shows the map calculated by subtracting the shorter from the longer timepoint. The chemical structures for the intermediates modeled at each timepoint are shown in the top row. The featureless maps for the $F_o$(5s)-$F_o$(3s), $F_o$(15s)-$F_o$(10s), $F_o$(20s)-$F_o$(3s), $F_o$(20s)-$F_o$(5s), $F_o$(30s)-$F_o$(10s), and $F_o$(30s)-$F_o$(15s) pairs indicate that the species present at these timepoints are identical. Therefore, we propose that DJ-1 catalyzes nearly two turnovers in the crystal, with the hemithioacetal present at 3,5, and 20s and the L-lactoylcysteine present at 10,15, and 30s. The matrix is symmetric about the diagonal, so only one unique half is shown. These maps are similar to those calculated using Careless-processed data in Fig. S5.

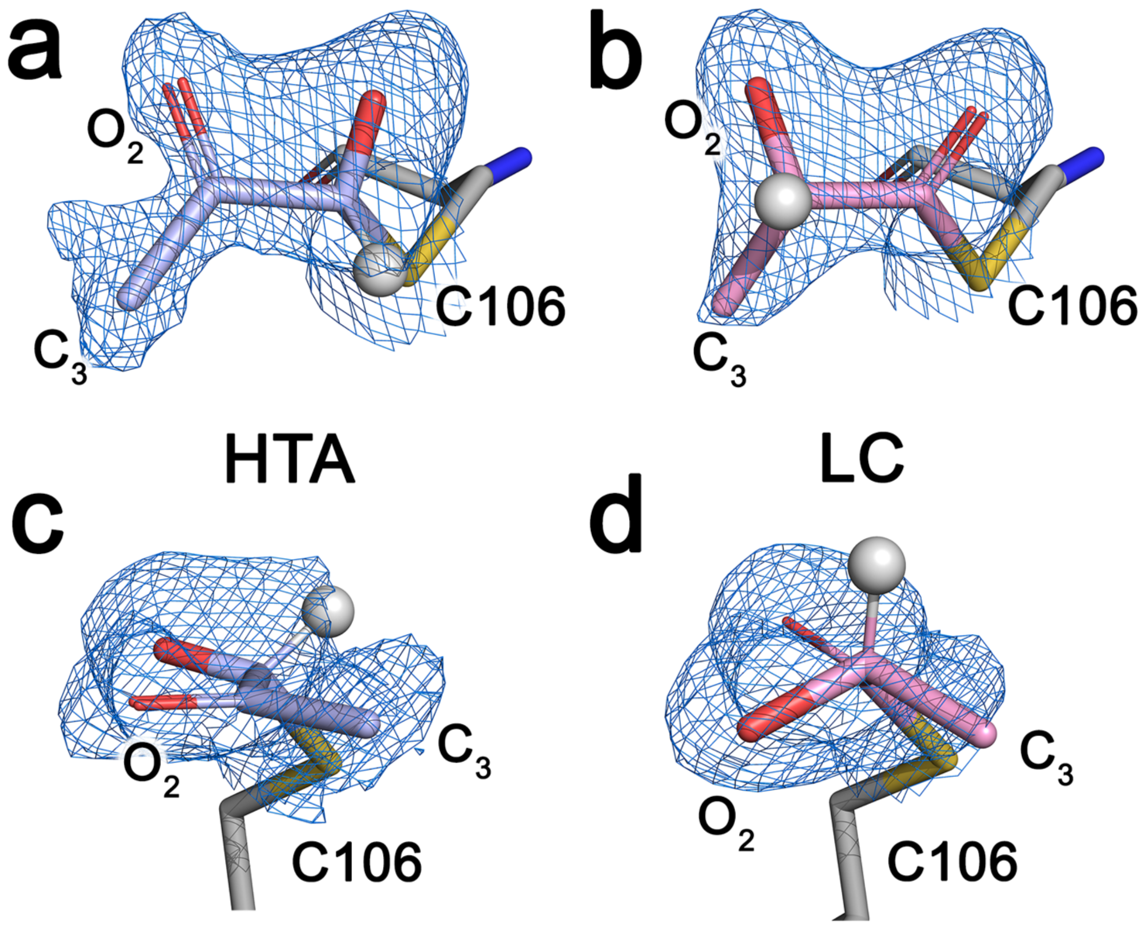


**Figure S7: Two views of methylglyoxal-hemithioacetal and L-lactoylcysteine in DJ-1 active site.** $2mF_o$-$DF_c$ electron density contoured at $1\sigma$ (blue) are shown for the hemithioacetal formed three seconds after mixing DJ-1 microcrystals with methylglyoxal (HTA, panels a and c) and the L-lactoylcysteine formed ten seconds after mixing DJ-1 microcrystals with methylglyoxal (LC, panels b and d). The key hydrogen atom whose position changes between the HTA and LC is shown as a white sphere and the Cys106 nucleophile is labeled. The more planar character of the HTA compared to LC is evident in the electron density, and the O2 oxygen and C3 carbon atoms (labeled) are more ordered in LC than in HTA, consistent with the $F_o$-$F_o$ electron density maps (Figs. S5 and S6).

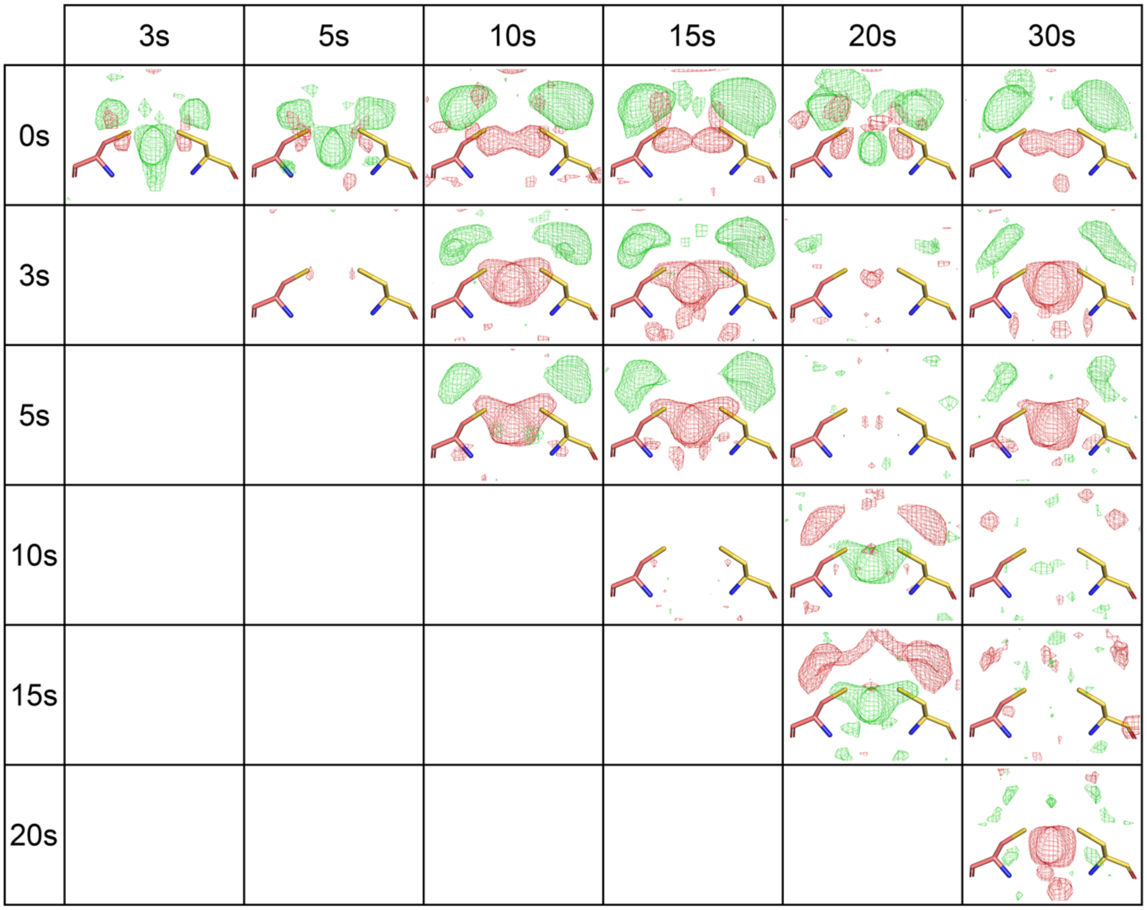


**Figure S8: Matrix of $F_o$-$F_o$ electron density maps at the Cys53 dimer interface, Careless-processed data.** $F_o$-$F_o$ electron density maps are contoured at +3σ (green) and -3σ (red) around the intermediate at Cys53 and its symmetry mate from the other protomer of the DJ-1 dimer (orange and yellow sticks). Each position in the matrix shows the map calculated by subtracting the shorter from the longer timepoint. As DJ-1 cycles through multiple catalytic cycles, strong difference features indicating motion of Cys53 appear, indicating allosteric communication between the active site and Cys53. Consistent with the $F_o$-$F_o$ electron density at the active site (Fig. S5), relatively featureless difference maps at the $F_o$(5s)-$F_o$(3s), $F_o$(15s)-$F_o$(10s), $F_o$(20s)-$F_o$(3s), $F_o$(20s)-$F_o$(5s), $F_o$(30s)-$F_o$(10s), and $F_o$(30s)-$F_o$(15s) timepoints indicate identical species.

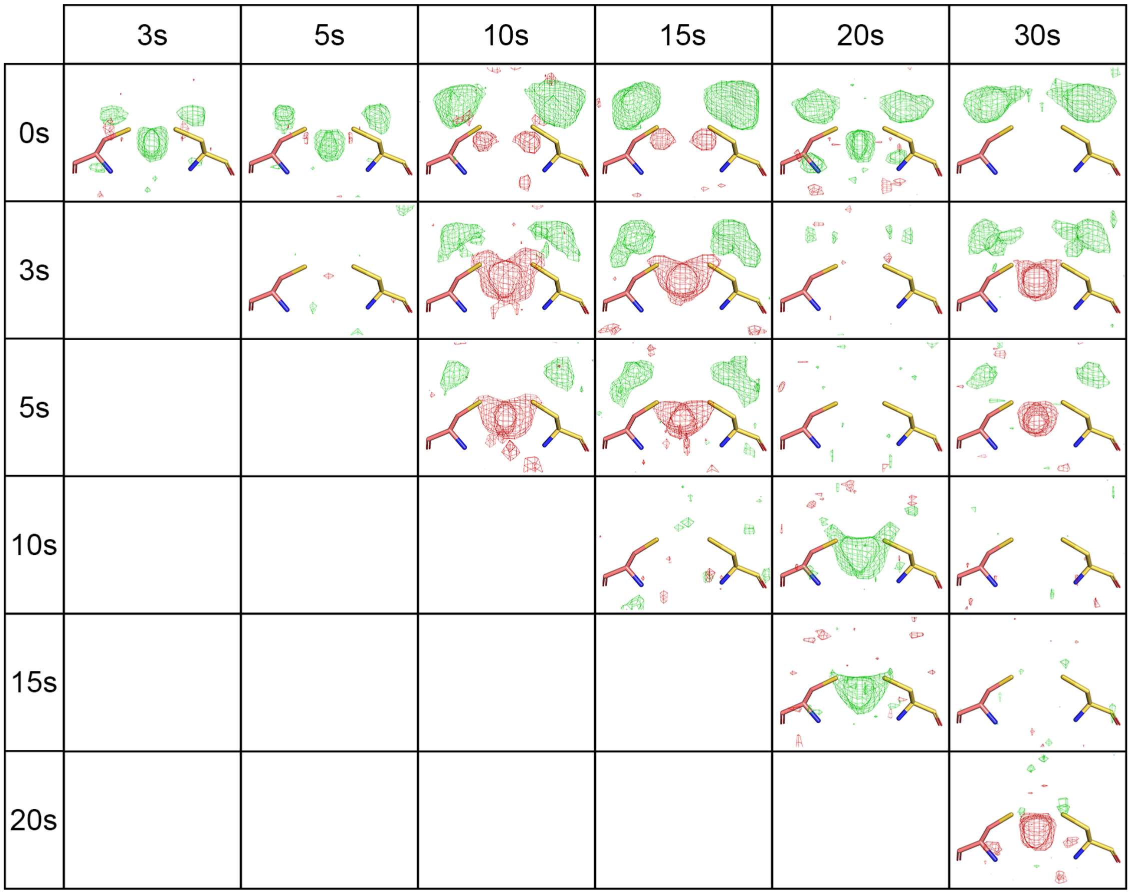


**Figure S9: Matrix of $F_o$-$F_o$ electron density maps at the Cys53 dimer interface, CrystFEL-processed data.** $F_o$-$F_o$ electron density maps are contoured at +3$\sigma$ (green) and -3$\sigma$ (red) around the intermediate at Cys53 and its symmetry mate from the other protomer of the DJ-1 dimer (orange and yellow sticks). Each position in the matrix shows the map calculated by subtracting the shorter from the longer timepoint. As DJ-1 cycles through multiple catalytic cycles, strong difference features indicating motion of Cys53 appear, indicating allosteric communication between the active site and Cys53. Consistent with the $F_o$-$F_o$ electron density at the active site (Fig. S5), relatively featureless difference maps at the $F_o$(5s)-$F_o$(3s), $F_o$(15s)-$F_o$(10s), $F_o$(20s)-$F_o$(3s), $F_o$(20s)-$F_o$(5s), $F_o$(30s)-$F_o$(10s), and $F_o$(30s)-$F_o$(15s) timepoints indicate identical species. These maps are similar to those calculated using Careless-processed data in Fig. S7.

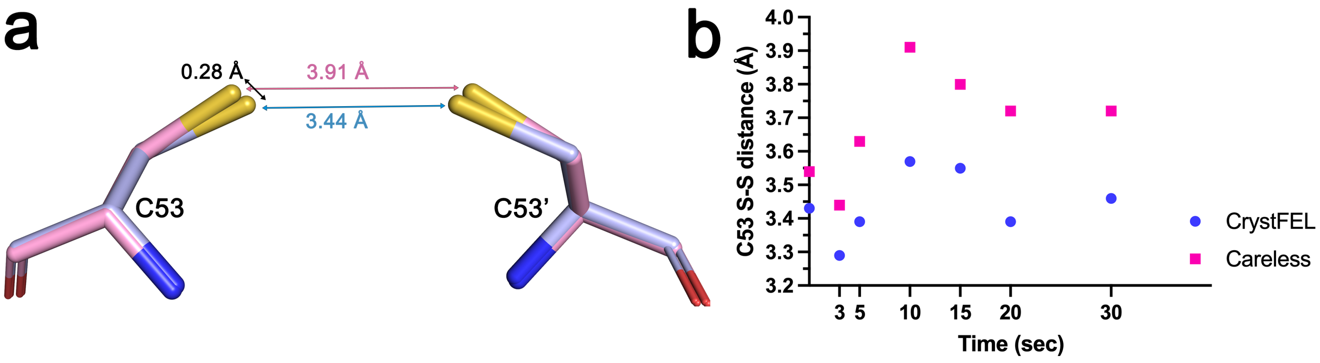


**Figure S10: Motion of Cys53 sidechain during catalysis.** Panel (a) shows the superposition of models at 3 sec (blue) and 10 sec (pink) after mixing with methylglyoxal refined against Careless-processed data. Even though these residues are 24 Å from the active site, the Cys53 Sγ atom moves by ~0.3 Å and the distance between the two symmetry-related Cys53 Sγ atoms contracts as DJ-1 transitions from the hemithioacetal species at 3 sec to the L-lactoylcysteine species at 10 sec. These motions result in the strong $F_o$-$F_o$ electron density features in Fig. S8 and S9. Panel (b) shows a plot of the Cys53-Cys53' Sγ distance as the reaction proceeds, showing a trend of shorter distances when the hemithioacetal is present (3,5,20 secs) and longer distances when the L-lactoylcysteine is present (10,15,30 sec). Distances are provided for models refined against CrystFEL-processed (blue circle) and Careless-processed (pink square) datasets, showing similar trends.

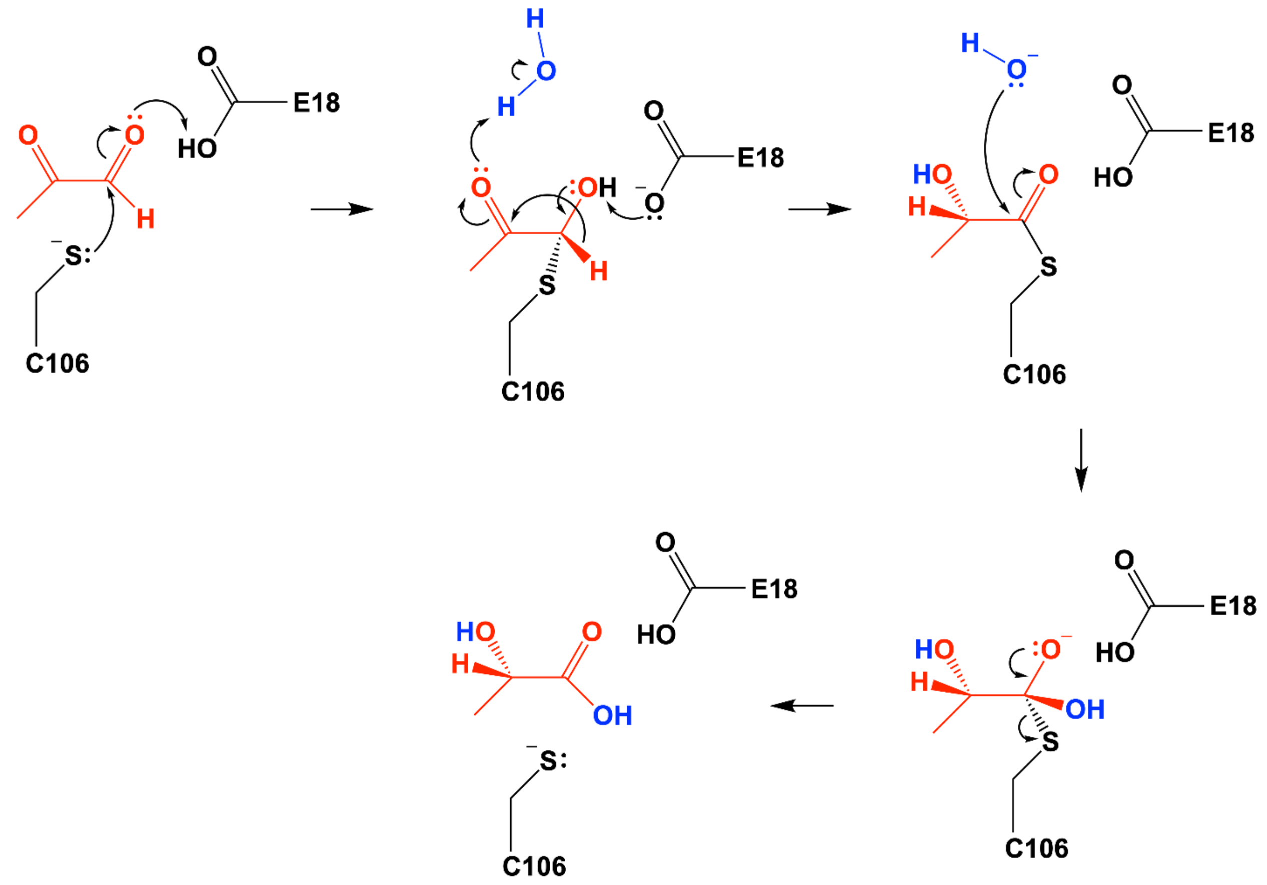


**Figure S11: Postulated alternative 1,2 hydride transfer mechanism for DJ-1 catalysis.** An alternative mechanism for concerted transfer of a hydride equivalent between the $C_1$ (aldehyde) and $C_2$ (ketone) atoms, reminiscent of an intramolecular Cannizzaro-like mechanism. The direction of electron flow is shown with curved arrows and colors indicate substrate-derived (red) or solvent-derived (blue) atoms.
